# METTL8 is required for 3-methylcytosine modification in human mitochondrial tRNAs

**DOI:** 10.1101/2021.05.02.442361

**Authors:** Jenna M. Lentini, Rachel Bargabos, Chen Chen, Dragony Fu

**Author notes:** Corresponding author: Dragony Fu, E–mail.

## Abstract

A subset of eukaryotic tRNAs is methylated in the anticodon loop to form the 3-methylcytosine (m3C) modification. In mammals, the number of tRNAs containing m3C has expanded to include mitochondrial (mt) tRNA-Ser-UGA and mt-tRNA-Thr-UGU. Whereas the enzymes catalyzing m3C formation in nuclear-encoded cytoplasmic tRNAs have been identified, the proteins responsible for m3C modification in mt-tRNAs are unknown. Here, we show that m3C formation in human mt-tRNAs is dependent upon the Methyltransferase-Like 8 (METTL8) enzyme. We find that METTL8 is a mitochondria-associated protein that interacts with mitochondrial seryl-tRNA synthetase along with mt-tRNAs containing m3C. Human cells deficient in METTL8 exhibit loss of m3C modification in mt-tRNAs but not nuclear-encoded tRNAs. Consistent with the mitochondrial import of METTL8, the formation of m3C in METTL8-deficient cells can be rescued by re-expression of wildtype METTL8 but not by a METTL8 variant lacking the N-terminal mitochondrial localization signal. Notably, METTL8-deficiency in human cells causes alterations in the native migration pattern of mt-tRNA-Ser-UGA suggesting a role for m3C in tRNA folding. Altogether, these findings demonstrate that METTL8 is required for m3C formation in mitochondrial tRNAs and uncover a potential role for m3C modification in mitochondrial tRNA structure.

## Introduction

The introduction of chemical modifications into RNA plays an important role in RNA processing, folding and function (Grosjean and Grosjean 2005). Transfer RNA (tRNA) is one of the most highly modified RNA species, accruing more than 10 modifications per tRNA on average (de Crecy-Lagard and Jaroch 2021; Pan 2018; Phizicky and Hopper 2015; Suzuki 2021). These diverse modifications can influence the folding and stability of tRNA species with subsequent impact on tRNA function in proper decoding and protein synthesis (Agris et al. 2017; El Yacoubi et al. 2012; Vare et al. 2017).

The human mitochondrial genome encodes 22 tRNA species that are essential for decoding the 13 mRNAs necessary for proper mitochondrial respiration (D’Souza and Minczuk 2018; Suzuki et al. 2011). Human mitochondrial tRNAs are transcribed by mitochondrial RNA polymerase and are subsequently modified by nuclear-encoded tRNA modification enzymes that are imported into the mitochondria (Putz et al. 2007; Suzuki and Suzuki 2014; Tiranti et al. 1997). Anticodon loop modifications in mt-tRNA play a critical role in mitochondrial protein synthesis by allowing the 22 mt-tRNAs increased decoding capacity for all 60 codons (Bohnsack and Sloan 2018). Moreover, modifications elsewhere in the sequence of mt-tRNAs are necessary for efficient folding and stability (Helm et al. 1998; Sakurai et al. 2005). Several human diseases are associated with pathogenic variants of mt-tRNA modification enzymes, underscoring the importance of mt-tRNA modifications in tRNA function and cellular physiology (Asano et al. 2018; Bohnsack and Sloan 2018; Kirino and Suzuki 2005; Lin et al. 2018; Pereira et al. 2018; Richter et al. 2018; Suzuki et al. 2011).

In human cells, mt-tRNA-Ser-UGA and mt-tRNA-Thr-UGU contain the 3-methylcytosine (m3C) modification at position 32 of the anticodon loop (Figure 1) (Cui et al. 2020; Suzuki and Suzuki 2014; Suzuki et al. 2020). In addition to mt-tRNA-Ser-UGA and mt-tRNA-Thr-UGU, the m3C modification is present in a subset of human nuclear-encoded cytoplasmic tRNAs (Arimbasseri et al. 2016a). The m3C modification is hypothesized to influence tRNA folding since the nucleotide at position 32 can form a non-canonical base-pair with residue 38 in certain tRNAs (Auffinger and Westhof 1999, 2001; Olejniczak and Uhlenbeck 2006). Moreover, loss of m3C in cytoplasmic tRNA-Ser of mouse stem cells causes alterations in gene expression and ribosome occupancy (Ignatova et al. 2020). However, the exact molecular role of m3C in tRNAs remains unknown. Moreover, the enzymes responsible for m3C formation in mitochondrial tRNAs have not been identified.

**Figure 1.**
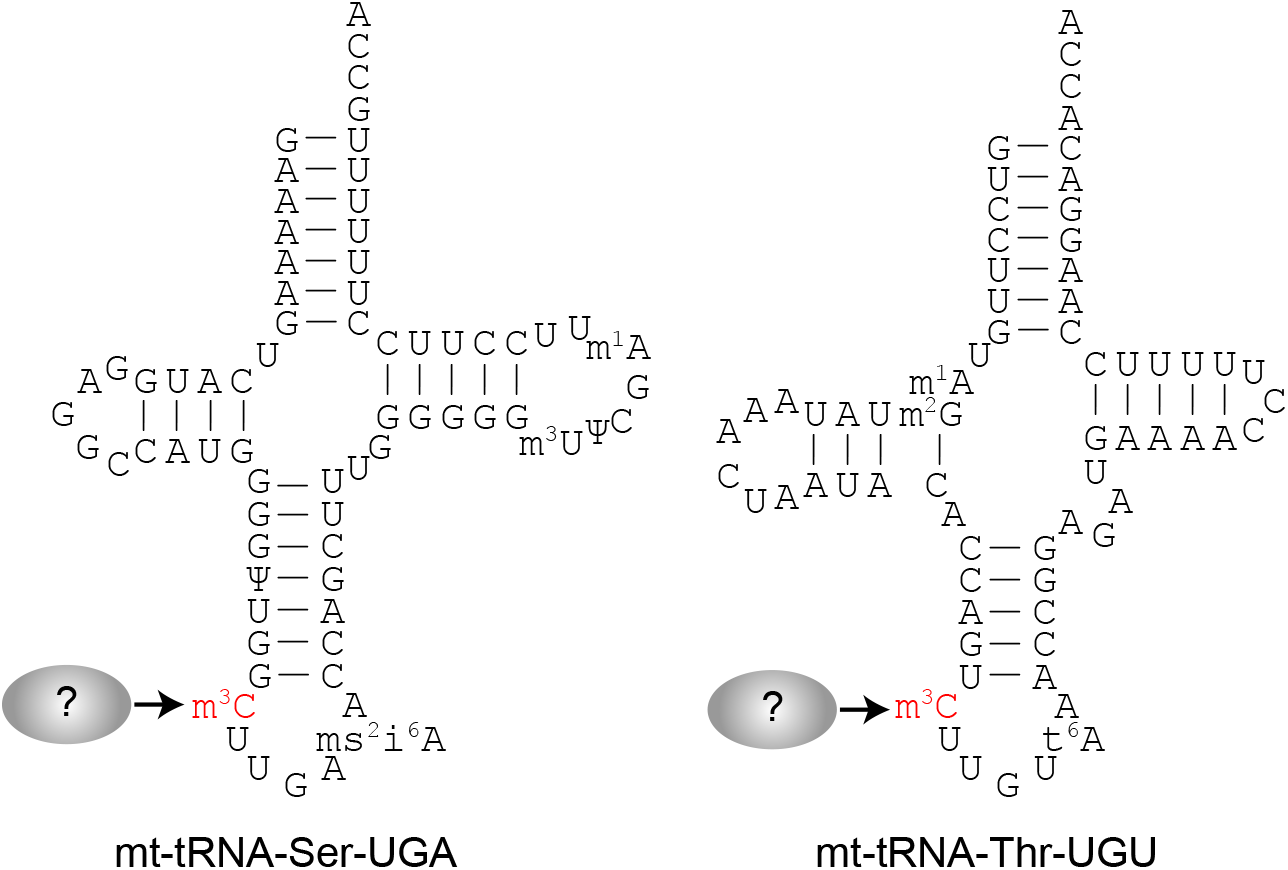
Secondary structures of human mitochondrial tRNAs containing the m3C modification. The m3C modification in the anticodon loop is shown in red.

In nuclear-encoded cytoplasmic tRNAs, the formation of m3C is dependent upon the Trm140p family of methyltransferases (D’Silva et al. 2011; Noma et al. 2011). While budding yeast *Saccharomyces cerevisiae* encodes a single Trm140 enzyme, fission yeast *Schizocaccharomyces pombe* encodes two Trm140 homologs that are separately responsible for modifying tRNA-Thr or tRNA-Ser isoacceptors (Arimbasseri et al. 2016a). Intriguingly, humans encode four Trm140 homologs that include METTL2A, METTL2B, METTL6, and METTL8. METTL2A and METTL2B encode paralogous proteins that share 99% amino acid sequence identity. Knockout of METTL2A/B in human cells abolishes m3C formation in a subset of cytoplasmic tRNA-Thr and tRNA-Arg isoacceptors while METTL6-deficiency in mouse cells leads to loss of m3C in cytoplasmic tRNA-Ser species (Xu et al. 2017). In addition, *S. cerevisiae* Trm140 forms an interaction with seryl-tRNA synthetase, which is required for stimulating the methyltransferase activity of Trm140p on tRNA-Ser isoacceptors (Han et al. 2017). We have also shown that human METTL2A and METTL2B interact with a tRNA synthetase mimic that is required for m3C formation in tRNA-Arg-CCU and UCU in human cells (Lentini et al. 2020). These studies highlight the evolutionary expansion of Trm140 homologs and the diversification of their RNA substrates through interactions with additional protein factors.

In contrast to METTL2A/B and METTL6, loss of METTL8 expression in human or mouse cells had no detectable impact on m3C formation in nuclear-encoded cytoplasmic tRNAs (Xu et al. 2017). Instead, METTL8-deficient mouse and human cells exhibited a reduction in m3C modification in poly-A selected RNAs that were depleted of tRNAs and rRNAs. These results suggest a possible role for METTL8 in the formation of m3C in mRNAs but the identity of these RNAs is unknown. In addition, METTL8 forms a nuclear RNA-binding complex that associates with numerous RNAs that could also be targets of methylation, including ribosomal RNAs (Zhang et al. 2020b). However, a recent study using a newly-developed sequencing method identified m3C in tRNAs but not in other RNAs above the limit of detection (Cui et al. 2020). Thus, the exact RNA targets of METTL8 remain equivocal.

Here, we show that METTL8 contains an N-terminal mitochondrial targeting sequence (MTS) that is necessary for association with mitochondria. Moreover, we find that METTL8 interacts with mitochondrial seryl-tRNA synthetase as well as m3C-containing mt-tRNAs. Using CRISPR-mediated gene editing, we demonstrate that METTL8-knockout in human cells abolishes m3C modification in mt-tRNA-Ser-UGA and mt-tRNA-Thr-UGU. Notably, mt-tRNA-Ser-UGA lacking m3C exhibits altered migration on native gels indicative of perturbed folding. In total, these results identify METTL8 as the protein responsible for m3C formation in mitochondrial tRNAs and uncover a putative role for m3C in maintaining proper tRNA folding.

## RESULTS

### METTL8 harbors an N-terminal mitochondrial localization signal

To identify proteins responsible for m3C formation in mitochondrial tRNAs, we first examined whether any of the four human Trm140 homologs possess a putative mitochondrial targeting sequence (MTS). Using predictive algorithms for identifying an N-terminal MTS (Fukasawa et al. 2015), human METTL6, METTL2A and METTL2B exhibited less than 3% probability of containing an MTS whereas METTL8 displayed an 84% probability of containing an MTS (Figure 2A, MTS, highlighted sequence). Moreover, METTL8 was predicted to contain a consensus cleavage site for the mitochondrial processing peptidase as well as sequence recognition motifs for the TOM20 mitochondrial import machinery (Figure 2A, MPP, TOM20). We also note that METTL8 has been detected in purified mitochondria from human and mouse cells via high-throughput proteomic experiments (Rath et al. 2021).

**Figure 2.**
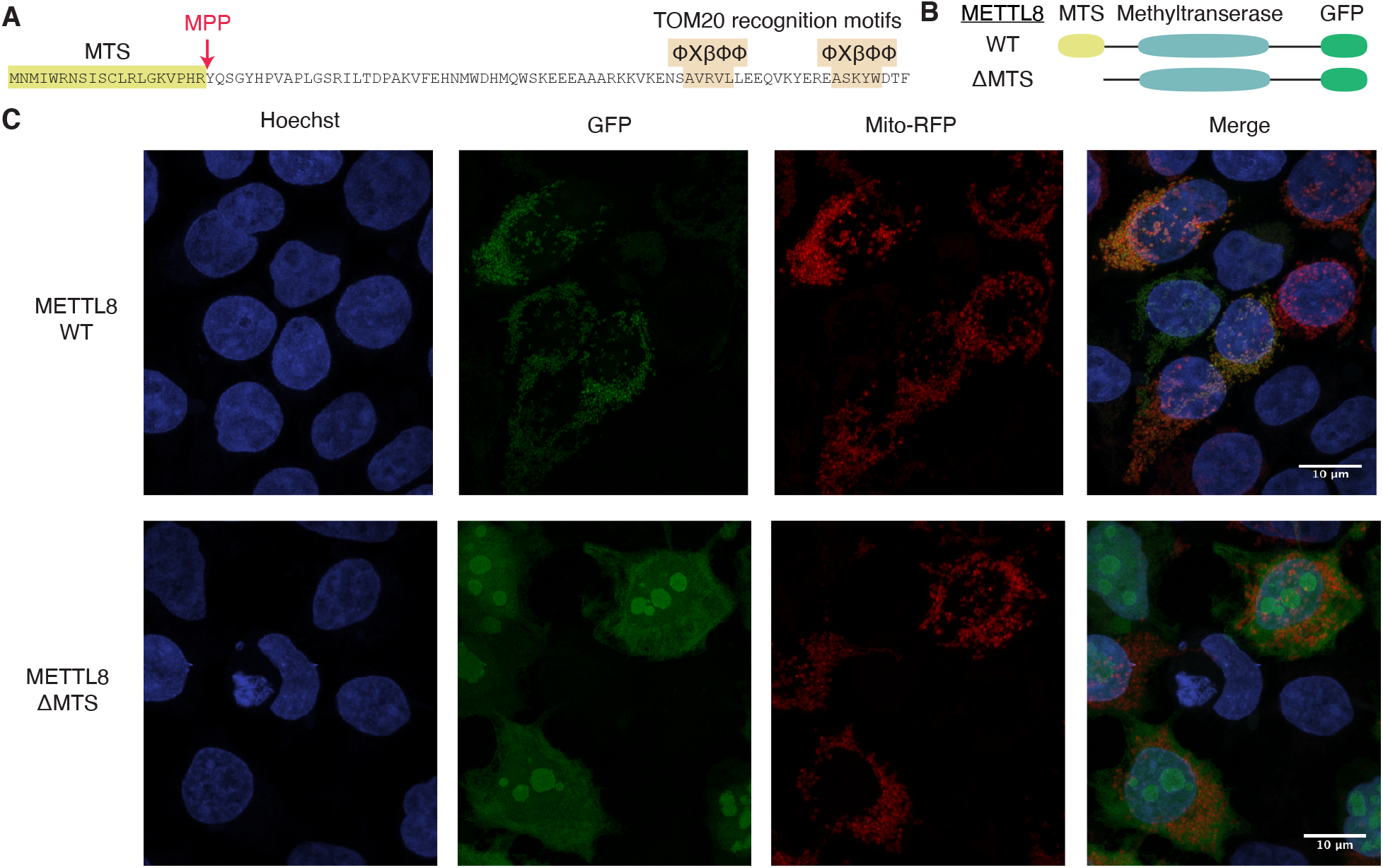
METTL8 exhibits localization in mitochondria that is dependent upon a mitochondrial targeting signal (MTS). (A) Sequence of the N-terminus of METTL8 with predicted MTS, proteolytic cleavage sites and TOM20 recognition motifs. ϕXβϕϕ is the TOM20 recognition motif where ϕ is a hydrophobic residue, X is any residue and β is a basic residue. (B) Representative schematic of METTL8 fusion proteins tagged with green fluorescent protein (GFP) at the carboxy terminus. Wildtype METTL8 or METTL8 lacking the MTS (ΔMTS) are depicted. (C) Confocal microscopy images of 293T cells transiently transfected with constructs expressing METTL8-WT or METTL8-ΔMTS fusion proteins with GFP. Mitochondria were identified using mitochondrion-targeted red fluorescent protein and nuclear DNA was stained with Hoechst. Overlap of red mitochondria and green GFP signal is displayed by yellow merged color.

To determine the subcellular localization of METTL8, we transiently transfected 293T human cells with constructs expressing human METTL8 fused at its carboxy terminus to green fluorescent protein (GFP) (Figure 2B, WT). We also tested the functional requirement for the predicted MTS in METTL8 by generating a construct expressing METTL8 lacking the MTS signal (Figure 2B, ΔMTS). As a marker for mitochondria, we co-expressed a red fluorescent protein (RFP) fused to the MTS of E1 alpha pyruvate dehydrogenase with the METTL8-GFP fusion proteins. METTL8-GFP exhibited a punctate, fragmented pattern of cytoplasmic localization surrounding the nucleus in human cells (Figure 2C, METTL8-WT, GFP). The fluorescence signal for METTL8 exhibited overlap with the RFP-tagged mitochondrial marker in nearly all cells that expressed both METTL8-GFP and the mitochondrial RFP marker (Figure 2C, Merge, yellow signal, see Supplemental Figure 1 for more image fields). In contrast to wildtype METTL8, METTL8-ΔMTS exhibited diffuse cytoplasmic localization along with enrichment in the nucleolus (Figure 2C, METTL8-ΔMTS, GFP). Compared to the METTL8-GFP subcellular localization pattern, the diffuse cytoplasmic ΔMTS-GFP signal exhibited weak overlap with the RFP mitochondrial maker (Figure 2C, METTL8-ΔMTS, Merge). These findings provide evidence that METTL8 contains a functional MTS sequence that is necessary for proper mitochondrial import. Moreover, loss of the MTS results in the nuclear import of METTL8 and apparent localization into the nucleolus.

### METTL8 forms a complex with nuclear proteins as well as mitochondrial SARS2

Previous studies have shown that *S. cerevisiae* Trm140 interacts with seryl-tRNA synthetase which stimulates the methyltransferase activity of Trm140 with tRNA-Ser isoacceptors (Han et al. 2017). Moreover, we have previously shown that human METTL2 forms a complex with the tRNA synthetase mimic DALRD3 to catalyze m3C formation in tRNA-Arg-CCU and tRNA-Arg-UCU (Lentini et al. 2020). To gain insight into the protein interaction network of human METTL8, we stably expressed METTL8 fused to the Twin-Strep tag at the C-terminus in 293T human embryonic cells (Schmidt et al. 2013). METTL8-Strep was affinity purified from whole cell extracts on streptactin resin, eluted with biotin and analyzed by silver staining. As a control for background contaminants, we performed a parallel purification from human cells integrated with an empty vector. Compared to the control, the METTL8 purification yielded a complex profile of interacting proteins ranging in size from 10 to 200 kDa (Figure 3A).

**Figure 3.**
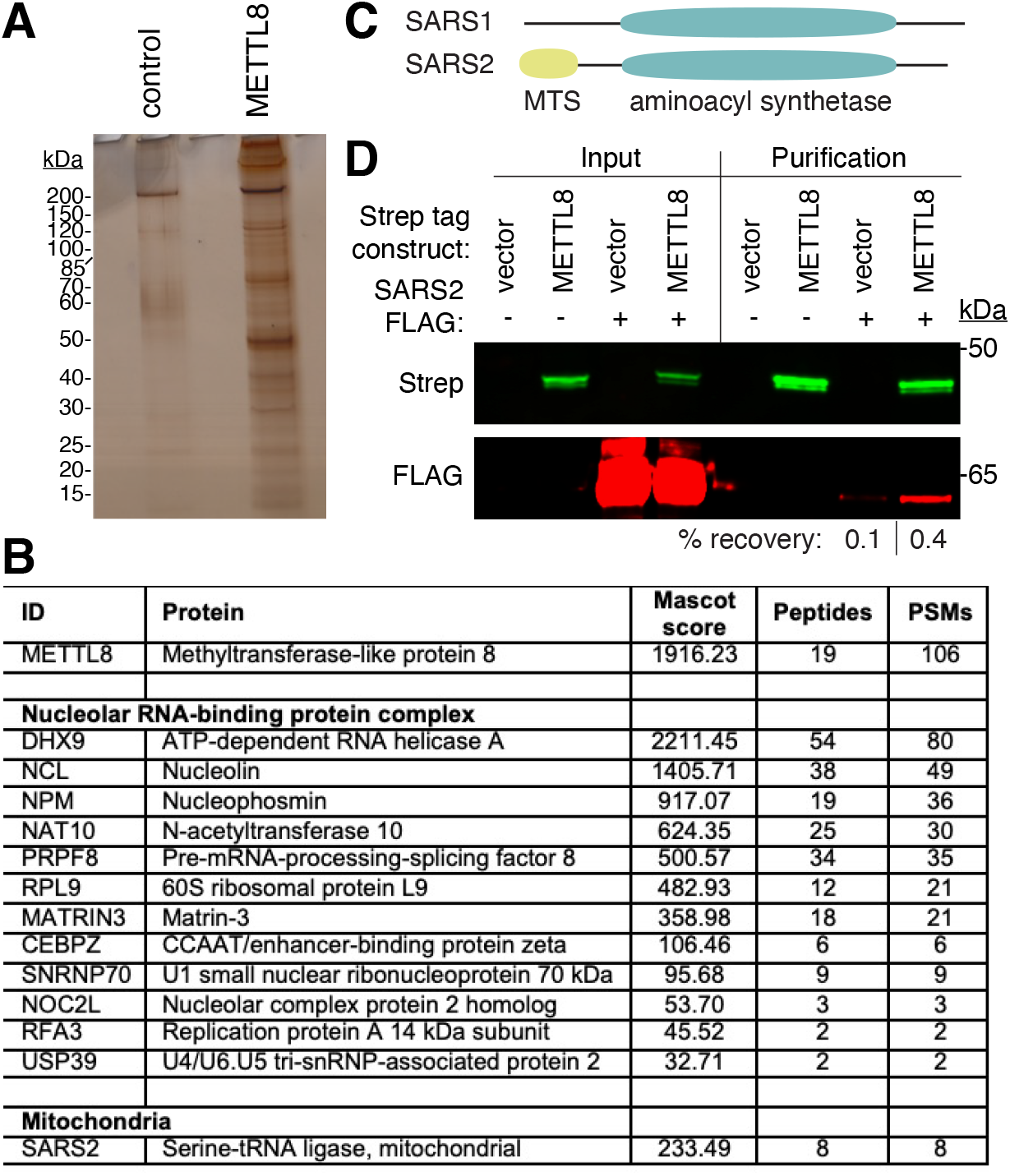
Identification of METTL8-interacting proteins. (A) Silver stain analysis of streptactin purifications from human 293T cell lines expressing either vector control or METTL8-Strep. (B) LC-MS analysis of peptides present in Strep-METTL8 purifications. Table lists a selection of proteins that were enriched in the Strep-METTL8 purification compared to empty vector control. Peptide spectral matches (PSM). (C) Schematic comparison of SARS1 versus SARS2 with aminoacyl synthetase domain and mitochondrial targeting signal (MTS) indicated. (D) Immunoblot analysis of empty vector control and Strep-METTL8 purifications from transient transfections. The immunoblot was probed with anti-Twin-Strep and anti-FLAG antibodies. Input represents 1% of total. % recovery represents the fraction of SARS2-FLAG recovered from the total input in the vector or METTL8 purification.

To identify METTL8-interacting proteins, we performed a large-scale purification from human cells expressing control or METTL8-Strep and subjected the entire eluted samples for proteomic analysis by liquid-chromatography mass spectrometry. After subtraction of contaminating proteins found in the control purification, we compiled an inventory of proteins that co-purified specifically with METTL8-Strep (Supplemental Data). Peptide sequences corresponding to METTL8 were detected in the purification from cells expressing METTL8-Strep, confirming the successful isolation of Strep-tagged METTL8 (Figure 3B, METTL8). Consistent with a previous study (Zhang et al. 2020a), we found that many METTL8-interacting proteins are part of a large nuclear complex enriched for proteins involved in RNA binding and R-loop regulation (Figure 3B, Nucleolar RNA-binding protein complex). Since the function of this nuclear METTL8 complex has been characterized, we focused our subsequent efforts on novel METTL8-interacting proteins.

In addition to nuclear proteins, we detected numerous peptides for mitochondrial seryl-tRNA synthetase (SARS2) in the METTL8 purification that were not found in the control purification (Figure 3B, SARS2). SARS2 shares a homologous aminoacyl synthetase domain as the cytoplasmic seryl-tRNA synthetase SARS1 but also contains an N-terminal mitochondrial targeting signal (MTS) (Figure 3C). No other tRNA synthetases were specifically found in the METTL8 purification. To confirm the interaction between METTL8 and mitochondrial SARS2, we transiently expressed METTL8-Strep with FLAG-tagged SARS2. Following purification of METTL8-Strep on streptactin resin, we monitored the copurification of SARS2-FLAG. Immunoblot analysis revealed the enrichment of SARS2-FLAG with Strep-METTL8 above the background in the vector control purification (Figure 3D). We note that SARS2 was not identified in a previous study on METTL8-interacting proteins, most likely due to the use of an N-terminal tag that would perturb localization of METTL8 to mitochondria (Zhang et al. 2020a). In contrast, we used a carboxy-terminal tag on METTL8 which would not disrupt recognition of the N-terminal MTS by the mitochondrial import machinery as shown with METTL8-GFP above. The identification of mitochondrial SARS2 as a METTL8-interacting protein suggests that METTL8 could play a role in modifying mitochondrial RNAs such as mt-tRNA-Ser-UGA, which contains m3C.

### METTL8 interacts with mitochondrial tRNAs

Based on the localization of METTL8 in mitochondria along with its interaction with mitochondrial SARS2, we next tested whether METTL8 displays stable interactions with mitochondrial tRNAs containing m3C. For these experiments, we transiently transfected 293T human cells with constructs expressing either METTL8-WT or METTL8-ΔMTS fused to the Strep tag. After purification on streptactin resin, a portion of each purification was retained for confirmation of protein isolation while the remainder was processed for RNA extraction (Figure 4A). Using immunoblotting, we confirmed the expression and purification of METTL8-WT or METTL8-ΔMTS on streptactin resin (Figure 4B). Both the METTL8-WT or METTL8-ΔMTS purifications exhibited an enrichment of 5S and 5.8S rRNA along with high molecular weight RNA species (Figure 4C, Supplemental Figure 2B). The purification of ribosomal RNA with METTL8 is consistent with the co-purification of ribosomal proteins along with the localization of METTL8 at R-loops in nucleolar rDNA genes (Zhang et al. 2020b).

**Figure 4.**
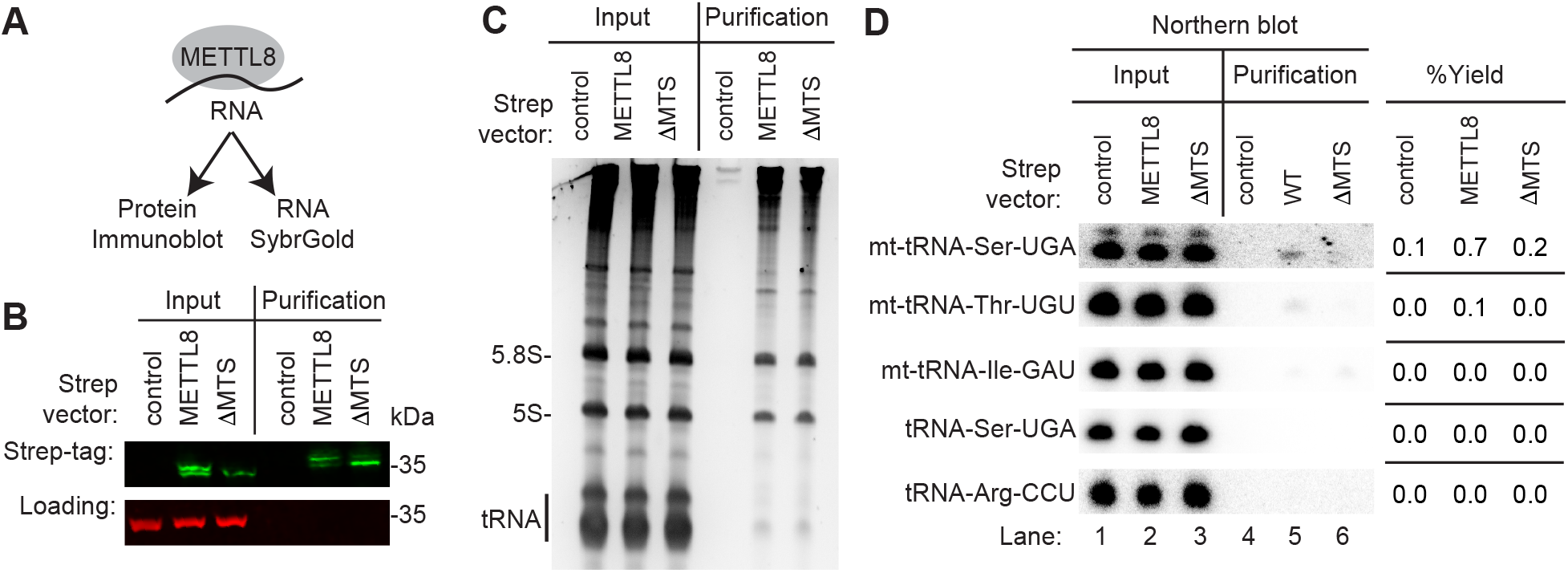
METTL8 interacts with mt-tRNA-Ser-UGA and mt-tRNA-Thr-UGU. (A) Purification of METTL8 and analysis of protein and RNA. (B) Immunoblot analysis of streptactin purifications from human cells expressing control, METTL8, or METTL8-ΔMTS fused to the twin-Strep tag. The immunoblot was probed with anti-TwinStrep and anti-actin antibodies. (C) Nucleic acid stain of RNAs extracted from the indicated input or purified samples after denaturing PAGE. The migration pattern of 5.8S rRNA (~150 nt), 5S rRNA (~120 nt) and tRNAs (~70–80 nt) are denoted. (D) Northern blot analysis of the gel in (C) using the indicated probes. Input represents 2% of total extracts used for purification. The percentage yield represents the amount of RNA in the Strep purification that was recovered from the total input. The experiment was repeated multiple times with comparable results (Supplemental Figure 2).

In addition to rRNA, we detected an RNA band migrating at the length of tRNAs that copurified with both METTL8 and METTL8-ΔMTS (Figure 4C). Based upon the mitochondrial localization of METTL8, we hypothesized that the tRNAs copurifying with METTL8 were the two mitochondrial tRNAs containing m3C; mt-tRNA-Ser-UGA and mt-tRNA-Thr-UGU. Using Northern blot probing, we found the METTL8 purification was enriched for mt-tRNA-Ser-UGA and mt-tRNA-Thr-UGU above the level of background binding in the control purification (Figure 4D, METTL8-WT). The enrichment of mt-tRNA-Ser-UGA and mt-tRNA-Thr-UGU was reduced in the METTL8-ΔMTS purification, consistent with the requirement for the MTS in METTL8 mitochondrial localization (Figure 4D, METTL8-ΔMTS). Moreover, neither METTL8 nor METTL8-ΔMTS exhibited enrichment above background for mt-tRNA-Ile, which lacks m3C (Figure 4D). In addition, no detectable enrichment was found for nuclear-encoded tRNAs containing m3C such as tRNA-Ser-UGA or tRNA-Arg-CCU in either METTL8 purification. The enrichment of mt-tRNA-Ser-UGA and mt-tRNA-Thr-UGU with METTL8 was repeated in an independent METTL8 purification with comparable results (Supplemental Figure 2). These results provide evidence that a subpopulation of METTL8 is imported into mitochondria where it interacts with mitochondrial tRNAs containing the m3C modification.

### METTL8 is required for m3C formation in mitochondrial tRNAs

The mitochondrial localization of METTL8 along with the copurification of mt-tRNA-Ser-UGA and mt-tRNA-Thr-UGU with METTL8 suggests that METTL8 could play a role in m3C formation in mitochondrial tRNAs. To investigate the functional role of METTL8, we generated human METTL8-knockout (KO) cell lines by CRISPR/Cas9 gene editing (Figure 5A). Using the HAP1 human haploid cell line, we generated two METTL8-KO cell clones containing a 72 base-pair deletion in exon 3 of the *METTL8* gene that was confirmed by PCR and Sanger sequencing (Figure 5B). The deletion is predicted to cause a translation frameshift that results in nonsense mediated decay or production of a truncated METTL8 missing the majority of the polypeptide. Using immunoblotting, we detected a substantial reduction of fulllength METTL8 protein in both METTL8-KO cell lines compared to the isogenic wildtype control cell lines (Figure 5C). No major change in morphology or proliferation were detected between the isogenic wildtype and METTL8-KO cell lines (data not shown).

**Figure 5.**
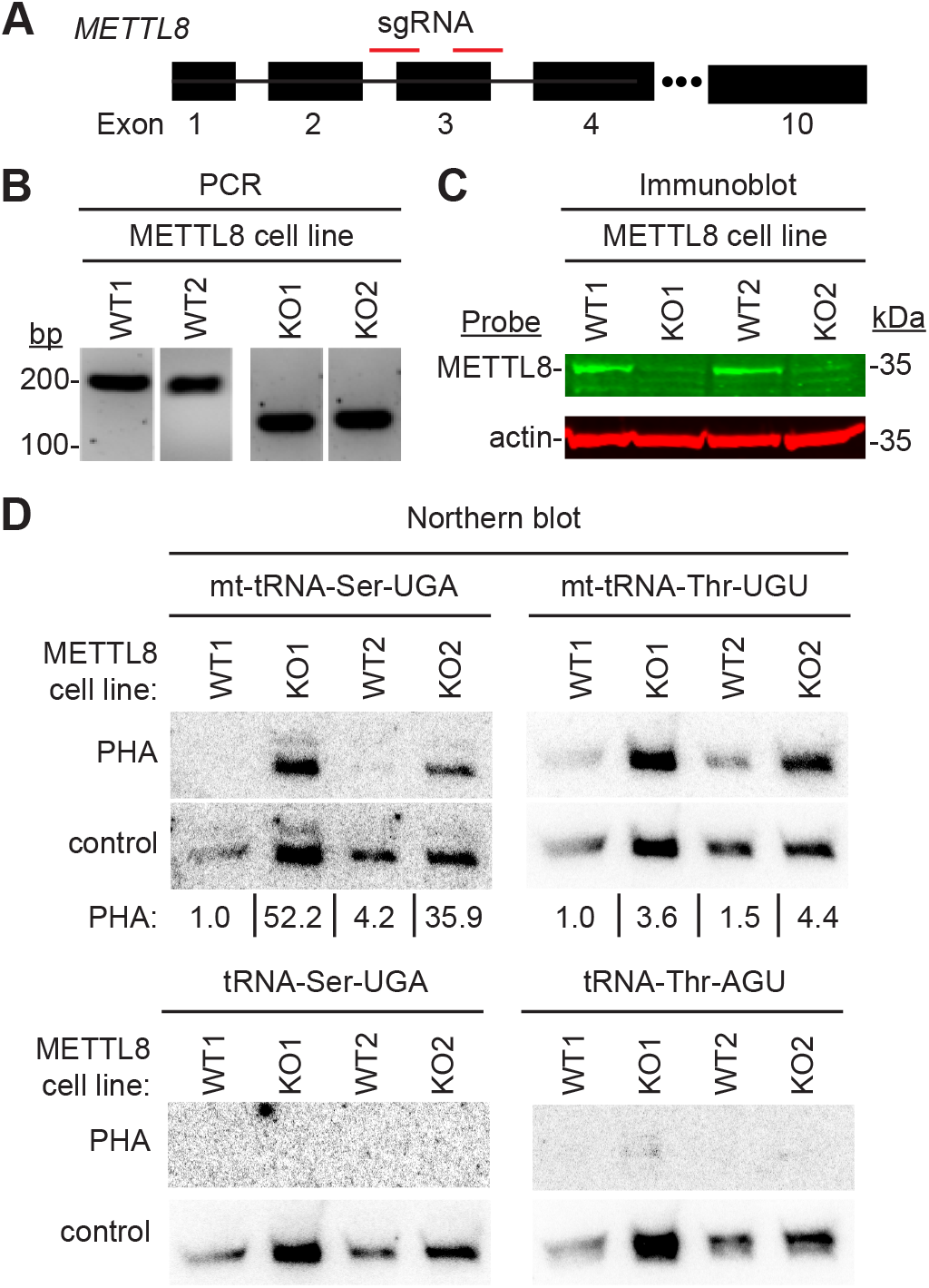
METTL8 is required for efficient m3C formation in mitochondrial tRNAs *in vivo*. (A) CRISPR/Cas9 gene knockout (KO) strategy depicting sequence guide RNAs (sgRNAs) targeting exon 3 of the human *METTL8* gene. (B) Genomic PCR demonstrating loss of exon 3 in METTL8-KO cell lines. Deletion of exon 3 results in a 72 base-pair deletion. (C) Immunoblot analysis of METTL8 expression in WT versus METTL8-KO human cell lines. Actin was used as a loading control. (D) Northern blot analysis using the Positive Hybridization in the Absence of Modification (PHA) assay with probes designed to detect m3C at position 32 and a control probe that hybridizes to a different area of the same tRNA. PHA quantification represents the ratio of PHA versus control probe signal expressed relative to the WT1 cell line. Northern blotting and quantification were performed on three independent cell preparations for each cell line with comparable results.

To monitor m3C formation in tRNA, we used the Positive Hybridization in the Absence of modification (PHA) assay (Arimbasseri et al. 2015; Dewe et al. 2017; Lamichhane et al. 2016; Lamichhane et al. 2011). This Northern blot-based assay relies on differential probe hybridization to tRNA caused by the presence or absence of m3C, which impairs base-pairing. Thus, a decrease in m3C modification leads to an increase in PHA probe signal that can be normalized against the probe signal from a different region of the same tRNA as an internal control. For mt-tRNA-Ser-UGA and mt-tRNA-Thr-UGU, there was a considerable increase in PHA probe signal in the METTL8-KO cell lines compared to WT, indicating the loss of m3C modification in these particular tRNAs (Figure 5D, mt-tRNA-Ser-UGA and mt-tRNA-Thr-UGU). The increase in PHA signal for mt-tRNA-Ser-UGA and mt-tRNA-Thr-UGU in the METTL8-KO cell lines was observed in independent probing experiments (Supplemental Figure 3). In contrast, no change in PHA signal was detected for nuclear-encoded tRNAs containing m3C in the METTL8-KO versus control cell lines (Figure 5D, tRNA-Ser-AGA and tRNA-Thr-AGU). We also note that the steady-state levels of all tested tRNAs were similar between the WT and METTL8-KO cell lines. These results suggest that METTL8 is required for the formation of m3C at position 32 of mitochondrial tRNA-Ser and mt-tRNA-Thr but not the other m3C-contaning tRNAs encoded by the nuclear genome.

To confirm that loss of m3C in mt-tRNAs is due to METTL8-deficiency, we tested whether re-expression of METTL8 in the METTL8-KO cell lines could rescue m3C formation in mt-tRNAs. In addition, we tested whether mitochondrial localization of METTL8 was necessary for m3C formation in mt-tRNAs using the METTL8-ΔMTS variant. We generated METTL8-KO cell lines containing an integrated lentiviral construct encoding the cDNAs for METTL8 or METTL8-ΔMTS fused to the Strep tag (Figure 6A). Due to the low expression level of the integrated METTL8 transgene in HAP1 human cells, we were unable to detect expression of METTL8 in cell lysates despite multiple attempts using the anti-METTL8 antibody (Supplemental Figure 4). As an alternative, we enriched for the METTL8-Strep fusion protein from cell lysates via purification on streptactin resin followed by immunoblotting with an anti-Strep tag antibody. Using this approach, the expression of either Strep-tagged METTL8-WT or METTL8-ΔMTS could be detected in both METTL8-KO cell lines (Figure 6B).

**Figure 6.**
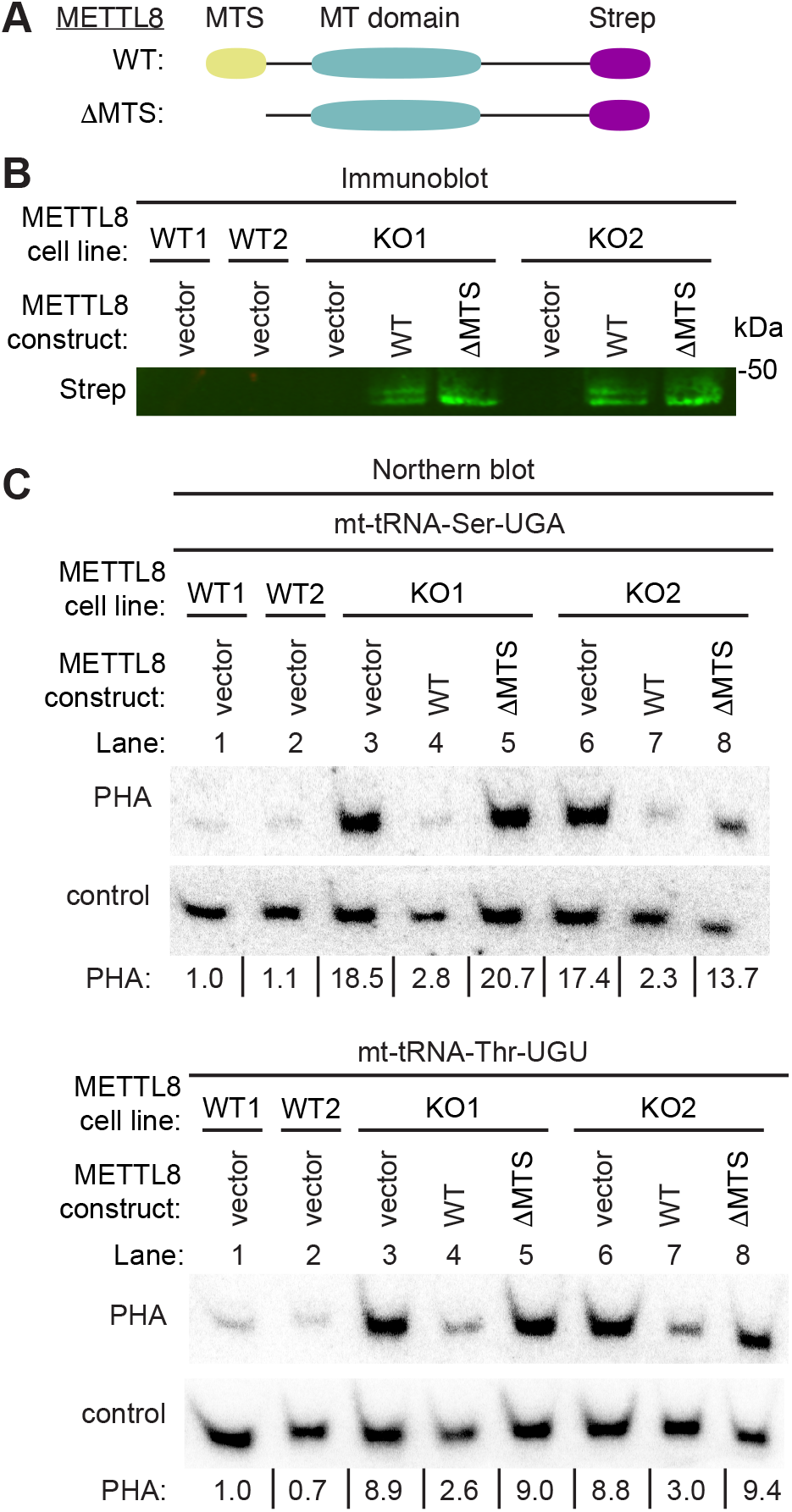
The MTS of METTL8 is required for efficient rescue of m3C formation in mitochondrial tRNAs of METTL8-KO human cells. (A) Schematic of METTL8 variants used for METTL8 rescue experiments. (B) Immunoblot analysis of Streptactin purifications from the indicated cell lines with anti-Strep antibodies. (C) Northern blot analysis using the Positive Hybridization in the Absence of Modification (PHA) assay with probes designed to detect m3C at position 32 and a control probe that hybridizes to a different area of the same tRNA. PHA quantification represents the ratio of PHA signal to control probe normalized to the WT1 cell line. (C) was repeated three times with similar results.

We next analyzed m3C status in the METTL8-KO cell lines expressing METTL8 or METTL8-ΔMTS. As expected, METTL8-KO cells with vector alone exhibited an increased PHA signal for mt-tRNA-Ser-UGA and mt-tRNA-Thr-UGU compared to WT cells indicative of m3C-deficiency (Figure 6C, compare lanes 1 and 2 to 3 and 6). Re-expression of METTL8-WT in either METTL8-KO cell line resulted in a reduced PHA signal for mt-tRNA-Ser-UGA and mt-tRNA-Thr-UGU compared to that of METTL8-KO cells with vector alone (Figure 6C, compare lanes 3 and 6 to lanes 4 and 7). The METTL8-KO cells reexpressing METTL8 exhibited comparable PHA signal for mt-tRNA-Ser-UGA and mt-tRNA-Thr-UGU to that of wildtype cells (Figure 6C, compare lanes 1 and 2 to lanes 4 and 7). In contrast to wildtype METTL8, expression of METTL8-ΔMTS had no major change in the PHA signal of mt-RNA-Ser-UGA or mt-tRNA-Thr-UGU in the METTL8-KO cell lines compared to vector alone (Figure 6C, compare lanes 3 and 6 to lanes 5 and 8). Overall, these results indicate that re-expression of METTL8 in METTL8-KO cells is sufficient to rescue m3C formation in mt-tRNAs and that mitochondrial localization is required for METTL8 function in m3C modification.

### m3C impacts the conformation of mitochondrial tRNA-Ser-UGA

The precise function of m3C modification in tRNA remains enigmatic. However, biophysical modeling studies suggest that the m3C modification could impact tRNA structure (Mao et al. 2021). To investigate a role for m3C in the folding of mt-tRNA-Ser-UGA and mt-tRNA-Thr-UGU, total RNA from WT or METTL8-KO human cells was resolved on non-denaturing native polyacrylamide gels followed by Northern blotting (Woodson and Koculi 2009). Both mt-tRNA-Ser-UGA and mt-tRNA-Thr-UGU from WT human cells migrated predominantly as a single band on native gels (Figure 7A, arrowhead). In contrast, we found that mt-tRNA-Ser-UGA from both METTL8-KO cell lines exhibited a shift to a slower-migrating species (Figure 7A, arrow). The slower-migrating mt-tRNA-Ser-UGA species in the METTL8-KO cell lines was detected in multiple independent native gel experiments (Supplemental Figure 5). The change in migration of mt-tRNA-Ser on native gels is likely due to conformation rather than length since no change in migration for mt-tRNA-Ser was detected on denaturing gels between WT versus METTL8-KO cells. Interestingly, no change in migration pattern was observed for mt-tRNA-Thr-UGU between WT versus METTL8-KO cell lines even though it also contains m3C dependent upon METTL8 (Figure 7A). We also observed no detectable change in migration of mitochondrial tRNA-Ile-GAU nor cytoplasmic tRNA-Arg-CCU between WT versus METTL8-KO cell lines.

**Figure 7.**
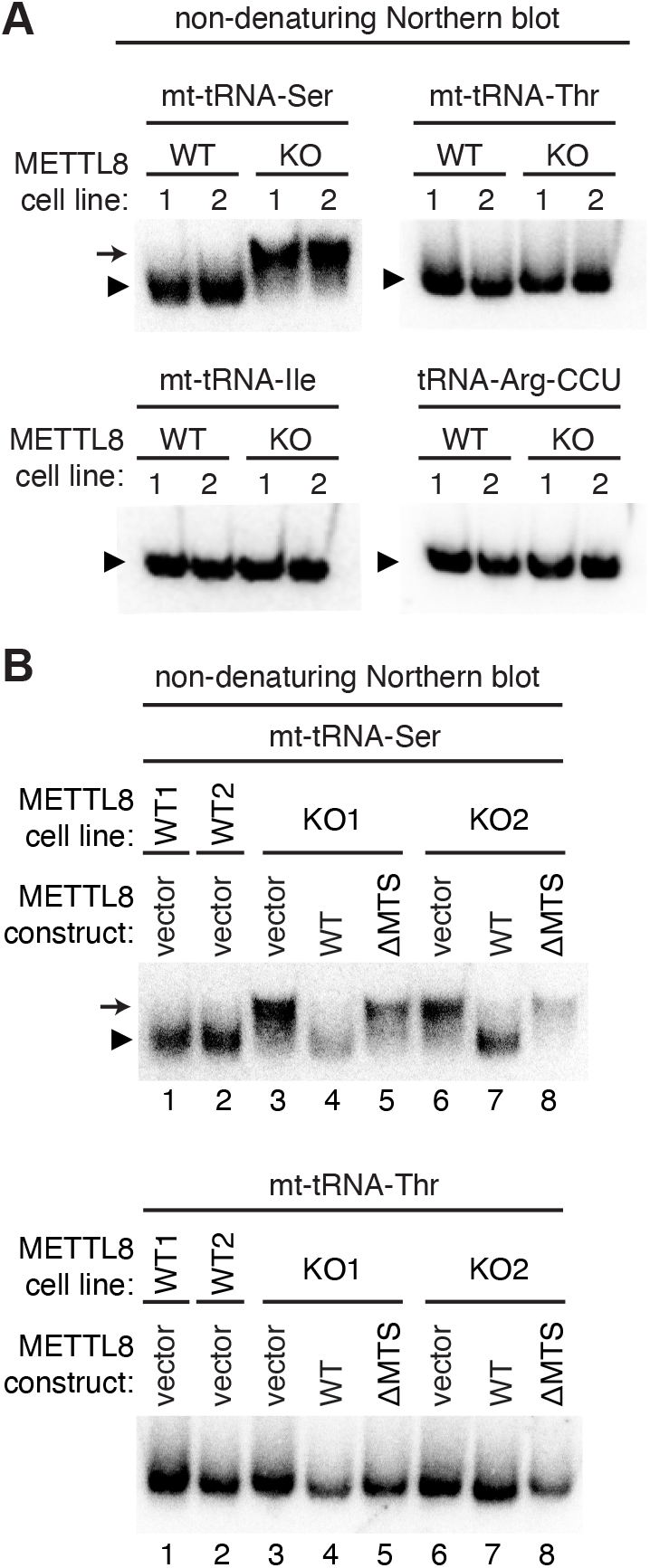
The native migration pattern of mt-tRNA-Ser is altered in METTL8-KO cell lines. (A) Total RNA from the indicated cell lines was fractionated on non-denaturing gels followed by transfer and hybridization with probes against the indicated tRNAs. The predominant band for each tRNA species in wildtype control cells is denoted by the arrowhead. The slower migrating mt-tRNA-Ser species found in METTL8-KO cell lines is denoted by the arrow. (B) Native gel analysis of mt-tRNA-Ser and Thr from the indicated cell lines. The predominant band for each tRNA species is denoted by the arrowhead and the slower migrating mt-tRNA-Ser species is denoted by the arrow.

To investigate the requirement for METTL8 in mt-tRNA-Ser structure, we examined native tRNA structure in the METTL8 cell lines expressing wildtype METTL8 or METTL8-ΔMTS. In METTL8-KO cell lines with vector alone, mt-tRNA-Ser-UGA existed predominantly as the slower migrating species compared to the faster-migrating species found in WT human cells with vector alone (Figure 7B, compare lanes 1 and 2 to lanes 3 and 6). Importantly, re-expression of METTL8 in the M8-KO cell line returned the migration pattern of mt-tRNA-Ser-UGA back to that of wildtype human cells consisting of predominantly the faster migrating species (Figure 7B, mt-tRNA-Ser, compare lanes 1 and 2 to lanes 4 and 7). In contrast, expression of METTL8-ΔMTS lacking mitochondrial localization was unable to restore the wildtype migration pattern of mt-tRNA-Ser-UGA (Figure 7B, compare lanes 1 and 2 to lanes 5 and 8). The mt-tRNA-Ser-UGA of METTL8-KO cell lines expressing METTL8-ΔMTS appeared more similar to the METTL8-KO cell lines with vector alone (Figure 7B, compare lanes 3 and 6 to lanes 5 and 8). Similar to the results above, METTL8-KO cell lines exhibited no detectable change in the migration pattern of mt-tRNA-Thr-UGU regardless of the integrated vector (Figure 7B, tRNA-Thr-UGU). Altogether, these findings suggest that m3C plays a role in mt-tRNA-Ser-UGA folding and conformation that is dependent upon the localization of METTL8 in mitochondria.

## Discussion

Here, we provide evidence that METTL8 is a nuclear-encoded tRNA modification enzyme responsible for catalyzing m3C formation in mt-tRNA-Ser-UGA and mt-tRNA-Thr-UGU. In addition, we show that METTL8 localization to the mitochondria is required for the efficient formation of m3C in mitochondrial tRNAs. Intriguingly, we find that METTL8-deficiency in human cells alters the mobility of mt-tRNA-Ser-UGA on native gels. To our knowledge, this provides the first experimental demonstration that m3C may impact the folding of an endogenous cellular tRNA.

Mitochondrial tRNAs fold into non-canonical tertiary structures that are inherently less stable (Watanabe et al. 2014). In particular, mt-tRNA-Ser-UGA exhibits a variant secondary structure with an extended anticodon stem that is hypothesized to maintain tertiary structure by compensating for additional changes elsewhere (Hanada et al. 2001; Yokogawa et al. 1991). Thus, m3C may also play an important role in facilitating the proper folding of the anticodon loop that impacts the overall tertiary structure of mt-tRNA-Ser-UGA. The results presented here suggest that mt-tRNA-Ser-UGA could exist in two different conformational states with m3C promoting the structure migrating as the slower-migrating form on native gels. The altered migration pattern of mt-tRNA-Ser-UGA could also be due to changes in modifications elsewhere in the tRNA that affects the overall conformation or charge state. The m3C modification could also play a role in the folding of mt-tRNA-Thr-UGU that might not be detectable using the native gel system. Future assays using structural probing could reveal the precise conformational differences observed between mitochondrial tRNAs with or without the m3C modification.

Deletion of Trm140p in combination with Trm1p in *S. cerevisiae* leads to a slight growth sensitivity to the translation inhibitor, cycloheximide (D’Silva et al. 2011). The lack of tRNA modifications catalyzed by Trm140 and Trm1 may impair the proper folding of tRNA substrates and their function in translation, thus rendering the yeast strain sensitive to perturbations in protein synthesis. While no major changes in the cytoplasmic polysome distribution profile was detected in METTL8-deficient human cells (Gu et al. 2018), our studies suggest that METTL8-catalyzed tRNA modification is likely to impact mitochondrial translation instead. Thus, it will be pertinent to assay for possible effects of m3C modification on mitochondrial protein synthesis in the future.

Using proteomics of purified METTL8 complexes, we have identified a novel interaction between METTL8 and mitochondrial seryl-tRNA synthetase (SARS2). This METTL8-SARS2 complex is analogous to the interaction previously found between *S. cerevisiae* Trm140 with seryl-tRNA synthetase and human METTL6 with cytoplasmic SARS1 (Han et al. 2017; Xu et al. 2017). Whereas seryl-tRNA synthetase has been shown to greatly stimulate the methyltransferase activity of Trm140, it is unknown whether human SARS1 or SARS2 are necessary for the methyltransferase activity of METTL6 or METTL8, respectively. Moreover, it is unclear how METTL8 specifically recognizes mt-tRNA-Ser-UGA but not mt-tRNA-Ser-AGY which also contains an unmodified C at the equivalent anticodon loop position but completely lacks a D-loop arm found in the canonical tRNA cloverleaf structure. Future studies using reconstitution of the METTL8-SARS2 complex *in vitro* could shed light on the role of SARS2 with METTL8 as well as the mechanism of tRNA recognition.

Intriguingly, METTL8 appears to have arisen within the vertebrate lineage (Arimbasseri et al. 2016b; D’Silva et al. 2011; Noma et al. 2011). Thus, METTL8 could have evolved distinct biological roles from the other Trm140 counterparts. Indeed, prior studies have linked METTL8 to adipogenesis as well as mouse embryonic stem cell differentiation (Badri et al. 2008; Gu et al. 2018; Jakkaraju et al. 2005). Moreover, METTL8 has also been found in a nucleolar RNA-binding complex that affects R-loop stability in genomic regions (Zhang et al. 2020b). However, the molecular role of METTL8 in these biological processes remains enigmatic. Our work suggests that METTL8 could play a key role in these diverse biological processes through the methylation of mitochondrial tRNAs and subsequent impact on mitochondrial physiology.

## Experimental Procedures

### Plasmids

The open reading frame (ORF) for METTL8 was RT-PCR amplified from HeLa human cervical carcinoma cell cDNA, cloned by restriction digest and verified by Sanger sequencing. The cDNA clones for METTL8 correspond to NM_001321154.2. METTL8 without the N-terminal MTS sequence was PCR amplified and cloned from the ORF of METTL8. Primers used are listed in Supplemental Table 1. The ORF for METTL8 and ΔMTS was cloned into either pcDNA3.1-Cterm-TWIN-Strep or pcDNA3.1-Cterm-GFP. The ORF for SARS2 was PCR amplified from cDNA plasmid HsCD00337767 (Plasmid Repository, Harvard Medical School) and cloned into pcDNA3.1-Cterm-3XFLAG.

For lentiviral expression constructs, Strep tagged-METTL8 and ΔMTS was first cloned into the pENTR.CMV.ON plasmid using NheI and NotI restriction sites. The entry vectors were recombined via LR clonase reaction (ThermoFisher) into the pLKO.DEST.puro destination vector to allow for stable integration of the target genes into human HAP1 cell lines.

### Mammalian tissue culture and CRISPR cell lines

The 293T human embryonic cell line was originally obtained from ATCC (CRL-3216). The HAP1 Human Male Chronic Myelogenous Leukemia (CML) cell line was obtained from Horizon Discovery Life Sciences. 293T human embryonic kidney cell lines were cultured in Dulbecco’s Minimal Essential Medium (DMEM) supplemented with 10% fetal bovine serum (FBS), 1X penicillin and streptomycin (ThermoFisher), and 1X Glutamax (Gibco) at 37 °C with 5% CO2. Cells were maintained and passaged every 3 days using 0.25% Trypsin. Human HAP1 AAVS wildtype and METTL8 knock-out cell lines were generated by CRISPR/Cas9 mutagenesis. Human HAP1 cell lines were cultured in Iscove’s Modified Dulbecco’s Medium (IMDM) supplemented with 10% FBS and 1X penicillin and streptomycin at 37 °C with 5% CO2. Cells were passaged every 3 days with 0.05% Trypsin.

For generation of stable cell lines, 2.5×10^5^ 293T cells were seeded onto 60 × 15mm tissue culture dishes. 1.25 μg of pLKO.DEST.puro plasmids containing the cloned ORF of METTL8-TWIN, ΔMTS-TWIN or empty vector along with a lentiviral packaging cocktail containing 0.75 μg of psPAX2 packaging plasmid and 0.5 μg of pMD2.G envelope plasmid was transfected into HEK293T cells using calcium phosphate transfection. Media was changed 16 hours post-transfection. 48 and 72 hours after transfection, media containing virus was collected and filter sterilized through 0.45 μm filters. 1 ml aliquots were flash frozen for later infection.

For lentiviral infection in 293T or HAP1 cell lines, 2.5×10^5^ cells were seeded in 6 well plates. 24 hours after initial seeding, 1 ml of either virus or media for mock infection along with 2 ml of media supplemented with 10 μg/ml of polybrene was added to each well. The cells were washed with PBS and fed fresh media 24 hours post-infection. Puromycin selection began 48 hours after infection at a concentration of 2 μg/ml. Fresh media supplemented with puromycin was added every other day and continued until the mock infection had no observable living cells. Proper integration and expression of each construct was verified via immunoblotting.

### Subcellular localization

The prediction of an N-terminal mitochondrial pre-sequence, cleavable localization signal and TOM20 import signal was performed using MitoFates: http://mitf.cbrc.jp/MitoFates/cgi-bin/top.cgi) (Fukasawa et al. 2015).

For localization of METTL8 tagged with GFP, 293T human cells were seeded onto coverslips in a 6-well plate followed by transfection with pcDNA3.1-METTL8-EGFP or pcDNA3.1-METTL8ΔMTS-EGFP using Lipofectamine 3000 reagent (Thermo Fisher). For mitochondrial localization, cells were infected with baculovirus expressing red fluorescent protein targeted to mitochondria (CellLight Mitochondria-RFP, BacMam 2.0, Life Technologies). To visualize the nucleus, Hoechst’s dye was added to the media for 30 minutes before cells were washed with PBS, fixed with 4% formaldehyde, and mounted in Aqua Poly/Mount (Polysciences Inc) followed by imaging on a Nikon A1R HD.

### Liquid chromatography-tandem mass spectrometry analysis

Vector control or METTL8-Strep stably integrated 293T cell lines were seeded on two 150 × 25 mm tissue culture dishes plates and grown until 70% confluency before cells of each line were combined and harvested for protein extraction. Cell pellets were resuspended in 1 mL of hypotonic lysis buffer (20 mM HEPES pH 7.9, 2 mM MgCl2, 0.2 mM EGTA, 10% glycerol, 0.1 mM PMSF, 1 mM DTT) per 150 × 25 mm tissue culture plate. Cells were kept on ice for 5 min and then underwent a freeze-thaw cycle three times to ensure proper detergent-independent cell lysis. NaCl was then added to the extracts at a concentration of 0.4 M and subsequently incubated on ice for 5 mins and spun down at 14,000 × g for 15 min at 4 °C. In all, 1 ml of Hypotonic Lysis buffer supplemented with 0.2% NP-40 was added to 500 μl of the supernatant extract.

Strep tagged proteins were then purified by incubating whole cell lysates with MagStrep “type3” XT beads (IBA Life Sciences) for five-six hours at 4 °C. Magnetic resin was washed three times in 20 mM HEPES pH 7.9, 2mM MgCl2, 0.2 mM EGTA, 10% glycerol, 0.1% NP-40, 0.2 M NaCl, 0.1 mM PMSF, and 1 mM DTT. Proteins were eluted with 1X Buffer BX (IBA LifeSciences) which contains 10 mM D-biotin. To ensure all protein was efficiently eluted off of the magnetic resin, the beads were left rotating in Buffer BX overnight at 4°C. Two one-hour elutions were completed the following day and all elutions were pooled together for each individual sample. The total eluate was then placed on a Spin-X UF 500 μl Centrifugal Concentrator (Corning) and spun at 15,000 × g for approximately 1 hour and 15 min at 4°C.

For protein separation, 15 μl of concentrated eluate was fractionated on a NuPAGE 4–12% Bis-Tris Protein gel (ThermoFisher). The gel was fixed overnight in 40% ethanol and 10% acetic acid. The gel was incubated in Sensitizing Solution (30% ethanol, 0.2% sodium thiosulphate and 6.8% sodium acetate) for 30 minutes before being washed three times with water for 5 min each wash. The gel was then stained in 0.25% silver nitrate for 20 min and washed twice more with water for 1 min each time. The bands were visualized by developing in 2.5% sodium carbonate and 0.015% formaldehyde and allowed to incubate until bands appeared. The remainder of each eluate (~65 μl) was loaded on a NuPAGE 4–12% Bis-Tris protein gel and briefly fractionated to yield a single gel band corresponding to the majority of proteins within each purification. Gel bands were excised using a razor blade and subject to in gel reduction, alkylation and trypsin digest by the URMC Mass Spectrometry Resource Lab.

Peptides were injected onto a homemade 30 cm C18 column with 1.8 μm beads (Sepax), with an Easy nLC-1000 HPLC (Thermo Fisher), connected to a Q Exactive Plus mass spectrometer (Thermo Fisher). Solvent A: 0.1% formic acid in water, Solvent B: 0.1% formic acid in acetonitrile. The gradient began at 3% B and held for 2 minutes, increased to 30% B over 13 minutes, increased to 70% over 2 minutes and held for 3 minutes, then returned to 3% B in 2 minutes and re-equilibrated for 8 minutes, for a total run time of 30 minutes. For peptides isolated from the single gel band corresponding to the majority of proteins, the gradient began at 3% B and held for 2 minutes, increased to 30% B over 41 minutes, increased to 70% over 3 minutes and held for 4 minutes, then returned to 3% B in 2 minutes and re-equilibrated for 8 minutes, for a total run time of 60 minutes. The Q Exactive Plus was operated in data-dependent mode, with a full MS1 scan followed by 8 data-dependent MS2 scans. The full scan was done over a range of 400-1400 m/z, with a resolution of 70,000 at m/z of 200, an AGC target of 1e6, and a maximum injection time of 50 ms. The MS2 scans were performed at 17,500 resolution, with an AGC target of 1e5 and a maximum injection time of 250 ms. The isolation width was 1.5 m/z, with an offset of 0.3 m/z, and a normalized collision energy of 27 was used.

Raw data was searched using the SEQUEST search engine within the Proteome Discoverer software platform, version 1.4 (Thermo Fisher), using the SwissProt human database that was downloaded in December of 2015. Trypsin was selected as the enzyme allowing up to 2 missed cleavages, with an MS1 mass tolerance of 10 ppm, and an MS2 mass tolerance of 25 mmu. Carbamidomethyl was set as a fixed modification, while oxidation of methionine was set as a variable modification. Percolator was used as the FDR calculator, filtering out peptides which had a q-value greater than 0.01(The et al. 2016). The Mascot scores of individual peptides were calculated as the absolute probability that the observed peptide match is a random event when matching spectra to all the expected spectra of a given proteome(Koenig et al. 2008). The Mascot score of a given peptide is equal to: −10 × Log_10_(P), where P is the absolute probability. The Mascot scores for individual proteins were then calculated based upon the summation of the individual peptides for all peptides matching a given protein. The LC-MS analysis was performed once with each sample.

### Transient Transfections and Protein-RNA Purifications

293T cells were transfected via calcium phosphate transfection method (Kingston et al. 2003). Briefly, 2.5×10^6^ cells were seeded on 100 × 20 mm tissue culture grade plates (Corning) followed by transfection with 10-20 μg of plasmid DNA. Media was exchanged 24 hours after transfection. Cells were harvested 48 hours post transfection by incubation in 0.25% trypsin and neutralization with media, followed by centrifugation of the cells at 700xg for 5 mins followed by subsequent PBS wash and a second centrifugation step.

Protein was extracted by the Hypotonic Lysis protocol immediately after cells were harvested posttransfection as was outlined in the above section. 0.5 ml of Hypotonic Lysis Buffer was added to each sample per 100 × 200 mm tissue culture plate, with an additional 0.5 ml of Hypotonic Lysis Buffer supplemented with NP-40 added for a final total of 1 ml of extract. Each extract was split into two 0.5 mL aliquots and flash frozen for later use. TWIN-Strep tagged proteins were then purified by incubating whole cell lysates from the transiently-transfected cell lines (0.5 ml) with 50 μl of MagStrep “type3” XT beads (IBA Life Sciences) for two-four hours at 4°C. Magnetic resin was washed three times in 20mM HEPES pH 7.9, 2 mM MgCl2, 0.2 mM EGTA, 10% glycerol, 0.1% NP-40, 0.2 M NaCl, 0.1 mM PMSF, and 1 mM DTT. Proteins were eluted with 1X Buffer BX (IBA LifeSciences). Purified proteins were visualized on a BOLT 4-12% Bis-Tris Plus gel (Life Technologies) and then transferred to Immobilon-FL Hydrophobic PVDF Transfer Membrane (Millipore Sigma) with subsequent immunoblotting against either the FLAG-epitope tag or TWIN-Strep epitope tag (Anti-FLAG M2, Sigma-Aldrich; THE^TM^ NWSHPQFEK antibody, GenScript).

We also utilized this purification procedure followed by TRIzol RNA extraction directly on the beads in order to identify co-purifying RNA with each protein of interest. Beads first underwent three washes in the Lysis Buffer as mentioned previously and then resuspended in 250 μl of Molecular Biology Grade RNAse-free water (Corning). 10 μl of the bead-water mixture was taken for immunoblotting analysis where the beads were mixed with 2X Laemmeli Sample Buffer (Bio-Rad) supplemented with DTT and boiled at 95°C for five minutes prior to loading onto a BOLT 4-12% Bis-Tris Plus gel (Life Technologies). RNA extraction followed the TRIzol LS RNA extraction protocol (Invitrogen). RNA was resuspended in 5 μl of RNAse-free water and loaded onto a 10% polyacrylamide, 7M urea gel. The gel was then stained with SYBR Gold nucleic acid stain (Invitrogen) to visualize RNA followed by transferring onto a Amersham Hybond-XL membrane and subsequent Northern blot procedures outlined below.

### CRISPR gene editing

CRISPR-Cas9 constructs were generated by cloning double-strand oligonucleotide inserts into pX333 (Addgene). sgRNAs were designed to target exon 3 of the METTL8 ORF (Table S1) and span a region of 72 bp. sgRNAs were also designed to target the AAVS ORF and act as our wildtype control. HAP1 human cells were seeded at 2.5×10^5^ in a 6-well plate and the next day transfected with px333-sgRNA4-sgRNA-5 using Lipofectamine 3000 (ThermoFischer). Single cell clones were isolated and placed into four 96-well plates. Single cell clones were allowed to grow until a single colony was visible. Media was replaced at the end of each week spanning around three weeks. Each single colony was trypsinized with 0.05% Trypsin and neutralized with media to re-seed in a single well of a 24-well plate and allowed again to grow. Once the wells reached ~70% confluency, cells were either harvested for DNA extraction (Qiagen) or frozen down for long-term storage. The presence of CRISPR-induced mutations in the *METTL8* gene was detected by PCR amplification and confirmed by Sanger sequencing using primers listed in Table S1.

### Immunoblot assays

To verify loss of METTL8, cell extracts were loaded onto BOLT 4–12% Bis-Tris gels (ThermoFisher) followed by immunoblotting onto PVDF membrane and probing with METTL8 antibody (ThermoFisher, cat. No PA557265, 1:500 dilution) and anti-Actin C4 (EMD Millipore, cat. No MAB1501, 1:000 dilution). Expression of METTL8 in stably-infected cell lines were characterized by immunoblotting with the anti-Strep antibody (THE^TM^ NWSHPQFEK antibody, Genscript, cat. No. A01732, 1:1000 dilution). Transient transfections of TWIN-Strep and FLAG-tagged proteins were also loaded onto BOLT 4–12% Bis-Tris gels, immunoblotted to PVDF membranes and probed with anti-TWIN-Strep (THE^TM^ NWSHPQFEK antibody, Genscript, cat. No. A01732, 1:1000 dilution) and anti-FLAG M2 (Anti-FLAG M2, Sigma-Aldrich, 1:3000 dilution). Image analysis of immunoblots were performed using Image Studio software (Li-Cor).

### RNA analysis

3-methylcytidine modification status was monitored using the Northern blot-based PHA assay. To conduct the PHA assay, probes were designed to hybridize upstream and downstream of residue 32 (oligos listed in Supplementary Table 1). A total of 5 μg of RNA was loaded onto a 10% polyacrylamide, 1xTBE, 7 M urea gel and transferred onto an Amersham Hybond-XL membrane (GE Healthcare) for Northern Blotting analysis. Oligonucleotides used to detect RNAs are listed in Supplementary Table 1. The oligos were radiolabeled by T4 polynucleotide kinase (NEB) with adenosine [γ32P]-triphosphate (6000 Ci/mmol, Amersham Biosciences) following standard procedures. Northern blots were visualized by Phosphor-Imager analysis and stripped via two incubations at 80 °C for 40 min in a buffer containing 0.15 M NaCl, 0.015 m Na-citrate and 0.1% SDS. Image analysis of Phosphorimager scans were performed using ImageJ open-source software.

For analysis of RNA structure under native conditions, 3 μg of RNA was resuspended in 6x Loading dye (Biolabs, cat. No B7024S) and separated on a 20% PA gel without urea run at 160 volts at 4°C. The RNA samples were kept on ice until loaded on the gel. RNA was then transferred onto an Amersham Hybond-XL membrane and subsequent probing with radiolabeled oligonucleotides as described above.

## Acknowledgements

We thank Eric Phizicky, Elaine Sia and Kejia Zhang for comments on the manuscript. We also thank Verda Kaye Thomas and the URMC Center for Advanced Light Microscopy and Nanoscopy for their assistance and resources for our microscopy imaging; Nicole Dawney for helping with microscopy analysis; and Kevin Welle and the URMC Mass Spectrometry Resource Lab for proteomic analysis.

---

**Supplemental Figure 1.**
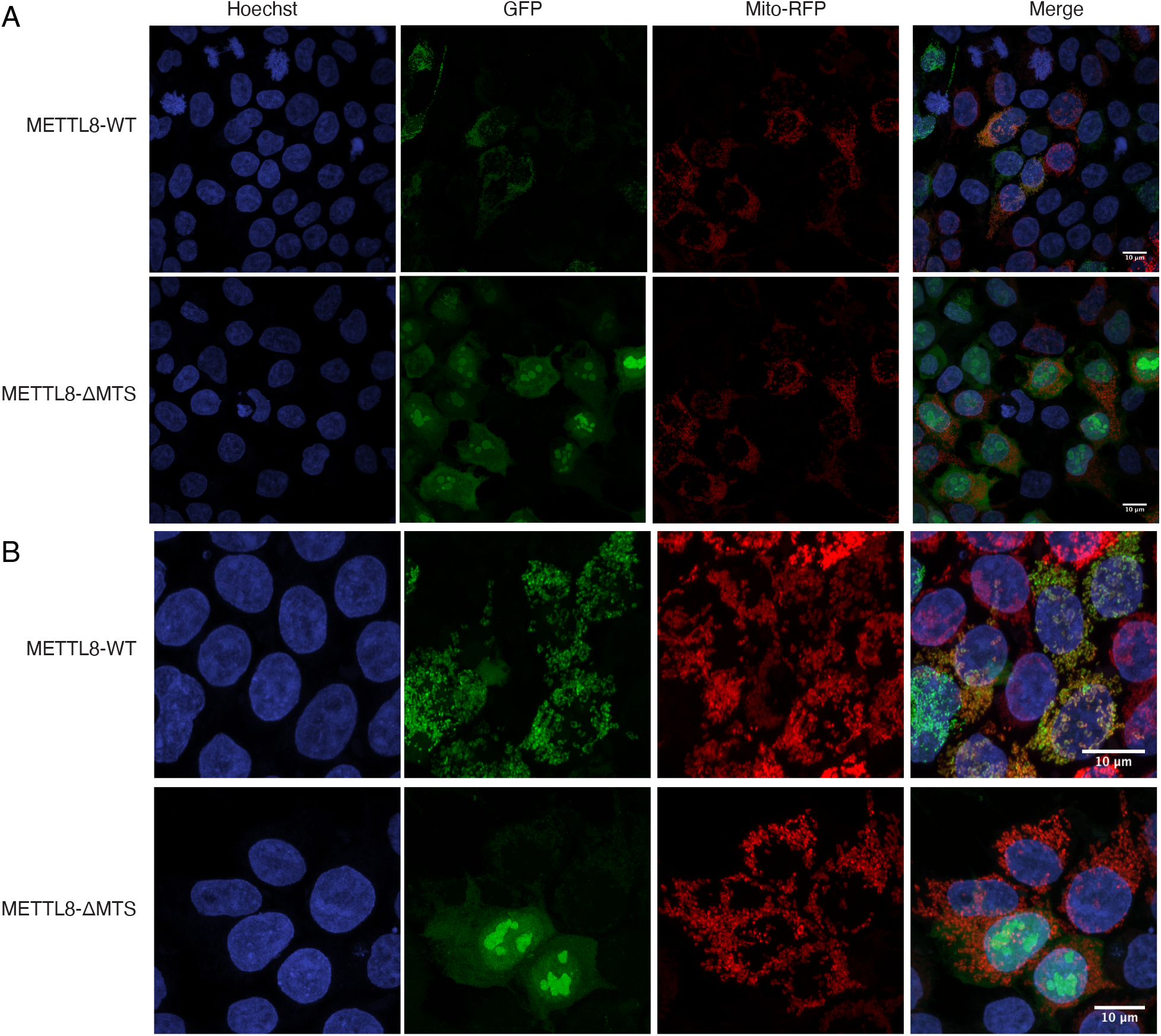
METTL8 exhibits localization in mitochondria that is dependent upon a mitochondrial targeting signal (MTS). (A, B) Confocal microscopy images of 293T cells transiently transfected with constructs expressing METTL8-WT or METTL8-ΔMTS fusion proteins with GFP. Mitochondria were identified using mitochondrion-targeted red fluorescent protein and nuclear DNA was stained with Hoechst. Overlap of red mitochondria and green GFP signal is displayed by yellow merged color.

**Supplemental Figure 2.**
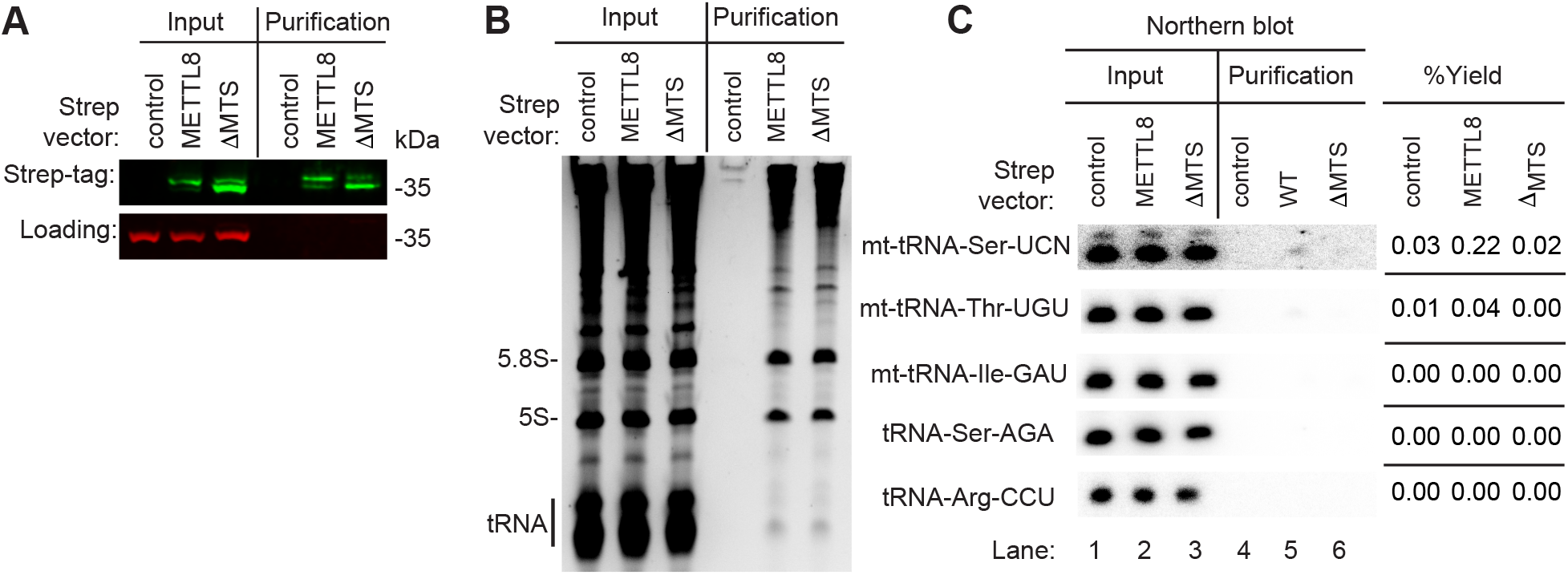
METTL8 interacts with mt-tRNA-Ser and mt-tRNA-Thr. Repeat of METTL8 purification in Figure 4. (A) Immunoblot analysis of streptactin purifications from human cells expressing control, METTL8, or METTL8-ΔMTS fused to the twin-Strep tag. The immunoblot was probed with anti-TwinStrep and anti-actin antibodies. (B) Nucleic acid stain of RNAs extracted from the indicated input or purified samples after denaturing PAGE. The migration pattern of 5.8S rRNA (~150 nt), 5S rRNA (~120 nt) and tRNAs (~70–80 nt) are denoted. (C) Northern blot analysis of the gel in (B) using the indicated probes. Input represents 2% of total extracts used for purification. The percentage yield represents the amount of RNA in the Strep purification that was recovered from the total input. Quantification was performed on the blot shown.

**Supplemental Figure 3.**
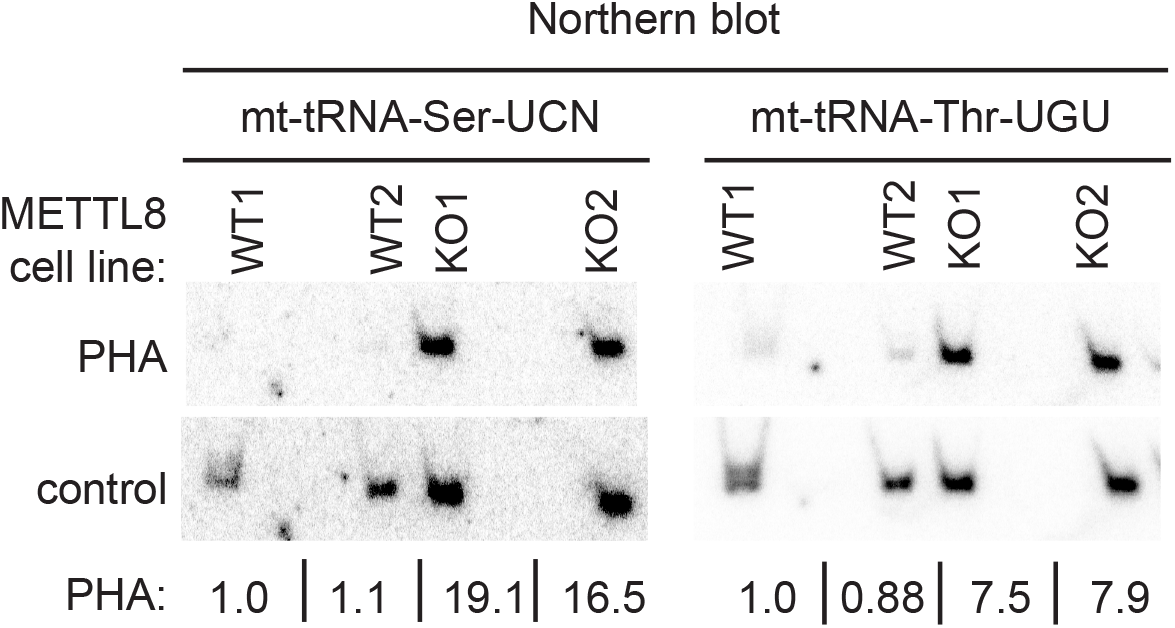
Repeat of Figure 5D. Northern blot analysis using the Positive Hybridization in the Absence of Modification (PHA) assay with probes designed to detect m3C at position 32 and a control probe that hybridizes to a different area of the same tRNA.

**Supplemental Figure 4.**
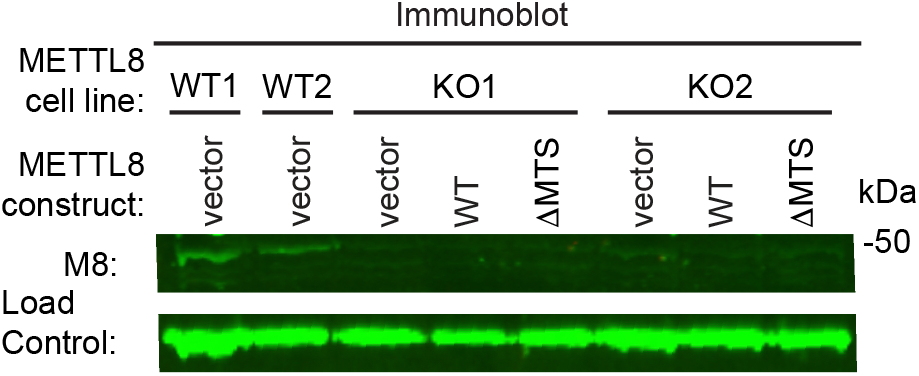
Immunoblot of the indicated human cell lines probed with anti-METTL8 antibody. Load Control is a non-specific band located on this gel.

**Supplemental Figure 5.**
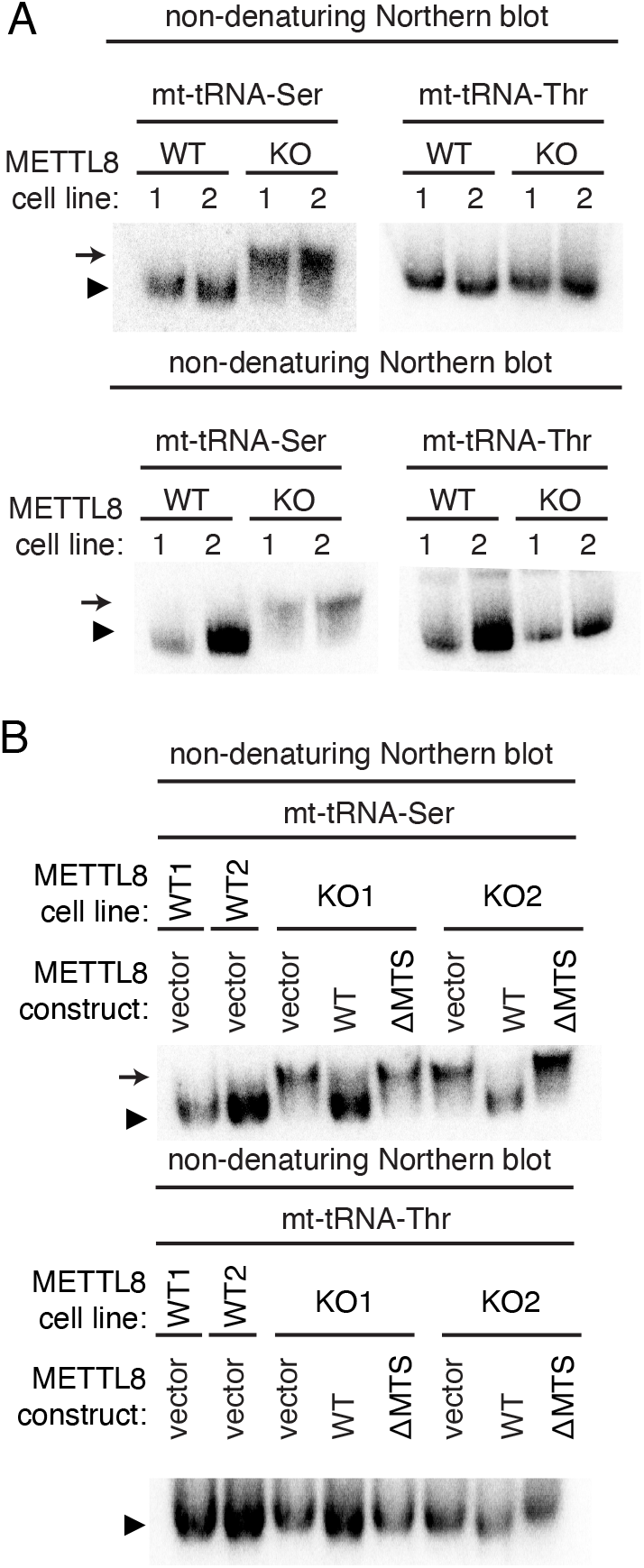
Independent replicate of Figure 7. The native migration pattern of mt-tRNA-Ser is altered in METTL8-KO cell lines. (A) Total RNA from the indicated cell lines was fractionated on nondenaturing gels followed by transfer and hybridization with probes against the indicated tRNAs. The predominant band for each tRNA species is denoted by the arrowhead. The slower migrating mt-tRNA-Ser species found in METTL8-KO cell lines is denoted by the arrow. (B) Native gel analysis of mt-tRNA-Ser and Thr from the indicated Rescue cell lines. The predominant band for each tRNA species is denoted by the arrowhead and the slower migrating mt-tRNA-Ser species is denoted by the arrow.

**Supplemental Table 1.**
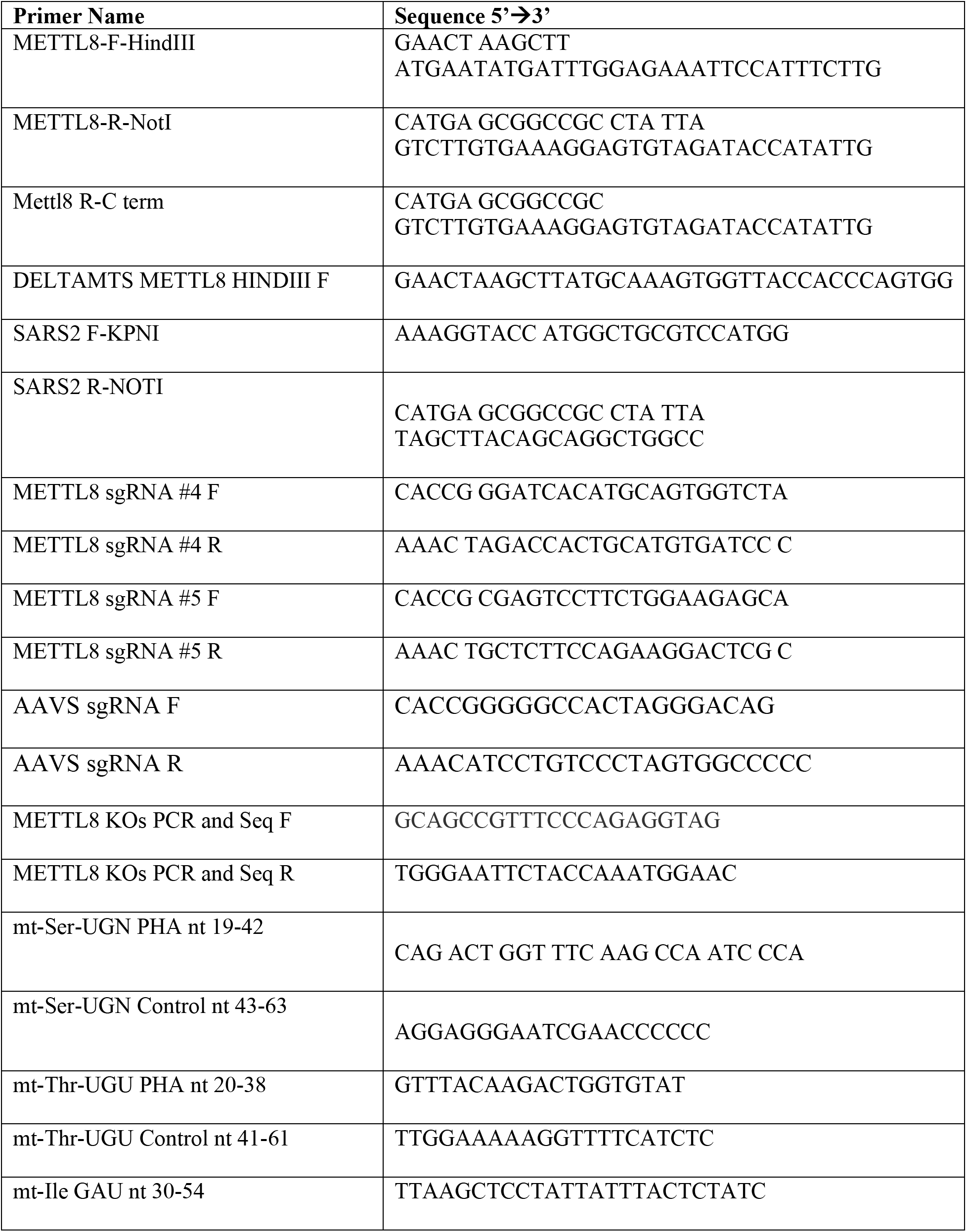
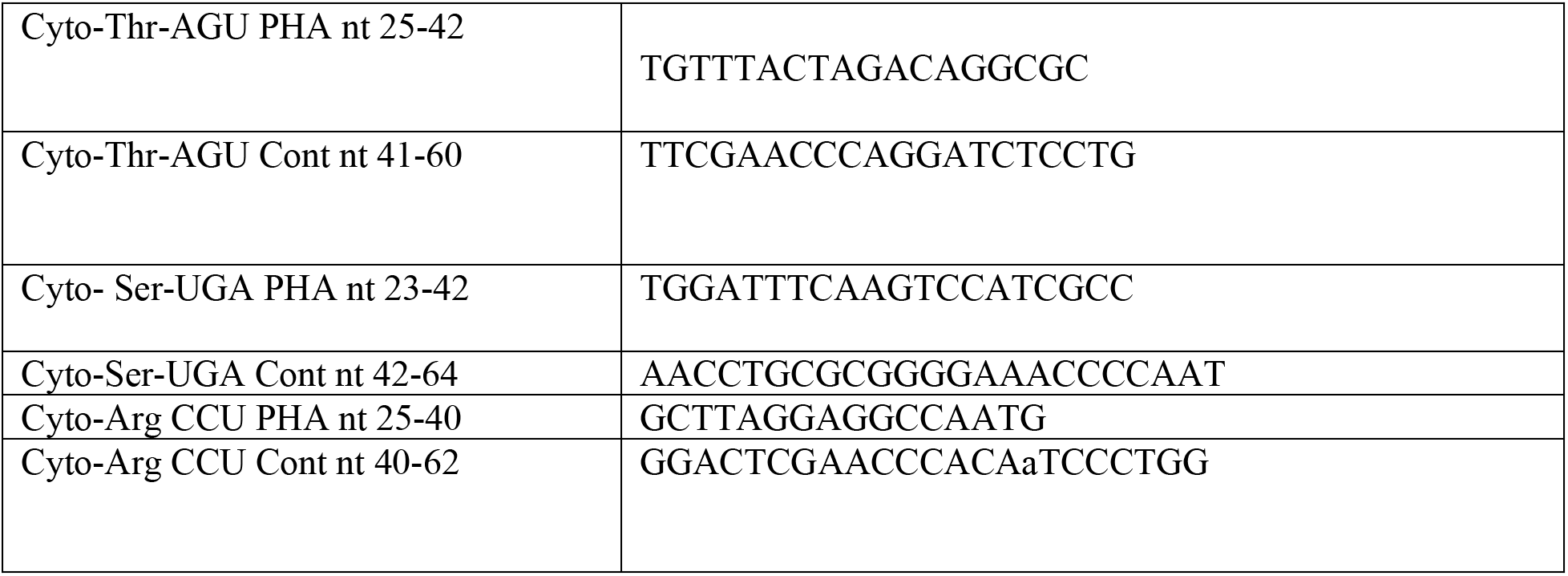
List of primers and oligonucleotides used in this study

## References

Agris PF, Narendran A, Sarachan K, Vare VYP, Eruysal E (2017) The Importance of Being Modified: The Role of RNA Modifications in Translational Fidelity. Enzymes 41: 1–50. doi: 10.1016/bs.enz.2017.03.005

Arimbasseri AG, Blewett NH, Iben JR, Lamichhane TN, Cherkasova V, Hafner M, Maraia RJ (2015) RNA Polymerase III Output Is Functionally Linked to tRNA Dimethyl-G26 Modification. PLoS Genet 11: e1005671. doi: 10.1371/journal.pgen.1005671

Arimbasseri AG, Iben J, Wei FY, Rijal K, Tomizawa K, Hafner M, Maraia RJ (2016a) Evolving specificity of tRNA 3-methyl-cytidine-32 (m3C32) modification: a subset of tRNAsSer requires N6-isopentenylation of A37. RNA. doi: 10.1261/rna.056259.116

Arimbasseri AG, Iben J, Wei FY, Rijal K, Tomizawa K, Hafner M, Maraia RJ (2016b) Evolving specificity of tRNA 3-methyl-cytidine-32 (m3C32) modification: a subset of tRNAsSer requires N6-isopentenylation of A37. RNA 22: 1400–10. doi: 10.1261/rna.056259.116

Asano K, Suzuki T, Saito A, Wei FY, Ikeuchi Y, Numata T, Tanaka R, Yamane Y, Yamamoto T, Goto T, Kishita Y, Murayama K, Ohtake A, Okazaki Y, Tomizawa K, Sakaguchi Y, Suzuki T (2018) Metabolic and chemical regulation of tRNA modification associated with taurine deficiency and human disease. Nucleic Acids Res 46: 1565–1583. doi: 10.1093/nar/gky068

Auffinger P, Westhof E (1999) Singly and bifurcated hydrogen-bonded base-pairs in tRNA anticodon hairpins and ribozymes. J Mol Biol 292: 467–83. doi: 10.1006/jmbi.1999.3080

Auffinger P, Westhof E (2001) An extended structural signature for the tRNA anticodon loop. RNA 7: 334–41. doi: 10.1017/s1355838201002382

Badri KR, Zhou Y, Dhru U, Aramgam S, Schuger L (2008) Effects of the SANT domain of tension-induced/inhibited proteins (TIPs), novel partners of the histone acetyltransferase p300, on p300 activity and TIP-6-induced adipogenesis. Mol Cell Biol 28: 6358–72. doi: 10.1128/MCB.00333-08

Bohnsack MT, Sloan KE (2018) The mitochondrial epitranscriptome: the roles of RNA modifications in mitochondrial translation and human disease. Cell Mol Life Sci 75: 241–260. doi: 10.1007/s00018-017-2598-6

Cui J, Liu Q, Sendinc E, Shi Y, Gregory RI (2020) Nucleotide resolution profiling of m3C RNA modification by HAC-seq. Nucleic Acids Res. doi: 10.1093/nar/gkaa1186

D’Silva S, Haider SJ, Phizicky EM (2011) A domain of the actin binding protein Abp140 is the yeast methyltransferase responsible for 3-methylcytidine modification in the tRNA anti-codon loop. RNA 17: 1100–10. doi: 10.1261/rna.2652611

D’Souza AR, Minczuk M (2018) Mitochondrial transcription and translation: overview. Essays Biochem 62: 309–320. doi: 10.1042/EBC20170102

de Crecy-Lagard V, Jaroch M (2021) Functions of Bacterial tRNA Modifications: From Ubiquity to Diversity. Trends Microbiol 29: 41–53. doi: 10.1016/j.tim.2020.06.010

Dewe JM, Fuller BL, Lentini JM, Kellner SM, Fu D (2017) TRMT1-Catalyzed tRNA Modifications Are Required for Redox Homeostasis To Ensure Proper Cellular Proliferation and Oxidative Stress Survival. Mol Cell Biol 37. doi: 10.1128/MCB.00214-17

El Yacoubi B, Bailly M, de Crecy-Lagard V (2012) Biosynthesis and function of posttranscriptional modifications of transfer RNAs. Annu Rev Genet 46: 69–95. doi: 10.1146/annurev-genet-110711-155641

Fukasawa Y, Tsuji J, Fu SC, Tomii K, Horton P, Imai K (2015) MitoFates: improved prediction of mitochondrial targeting sequences and their cleavage sites. Mol Cell Proteomics 14: 1113–26. doi: 10.1074/mcp.M114.043083

Grosjean H, Grosjean H (2005) Fine-tuning of RNA functions by modification and editing. Springer

Gu H, Do DV, Liu X, Xu L, Su Y, Nah JM, Wong Y, Li Y, Sheng N, Tilaye GA, Yang H, Guo H, Yan J, Fu XY (2018) The STAT3 Target Mettl8 Regulates Mouse ESC Differentiation via Inhibiting the JNK Pathway. Stem Cell Reports 10: 1807–1820. doi: 10.1016/j.stemcr.2018.03.022

Han L, Marcus E, D’Silva S, Phizicky EM (2017) S. cerevisiae Trm140 has two recognition modes for 3-methylcytidine modification of the anticodon loop of tRNA substrates. RNA 23: 406–419. doi: 10.1261/rna.059667.116

Hanada T, Suzuki T, Yokogawa T, Takemoto-Hori C, Sprinzl M, Watanabe K (2001) Translation ability of mitochondrial tRNAsSer with unusual secondary structures in an in vitro translation system of bovine mitochondria. Genes Cells 6: 1019–30. doi: 10.1046/j.1365-2443.2001.00491.x

Helm M, Brule H, Degoul F, Cepanec C, Leroux JP, Giege R, Florentz C (1998) The presence of modified nucleotides is required for cloverleaf folding of a human mitochondrial tRNA. Nucleic Acids Res 26: 1636–43.

Ignatova VV, Kaiser S, Ho JSY, Bing X, Stolz P, Tan YX, Lee CL, Gay FPH, Lastres PR, Gerlini R, Rathkolb B, Aguilar-Pimentel A, Sanz-Moreno A, Klein-Rodewald T, Calzada-Wack J, Ibragimov E, Valenta M, Lukauskas S, Pavesi A, Marschall S, Leuchtenberger S, Fuchs H, Gailus-Durner V, de Angelis MH, Bultmann S, Rando OJ, Guccione E, Kellner SM, Schneider R (2020) METTL6 is a tRNA m(3)C methyltransferase that regulates pluripotency and tumor cell growth. Sci Adv 6: eaaz4551. doi: 10.1126/sciadv.aaz4551

Jakkaraju S, Zhe X, Pan D, Choudhury R, Schuger L (2005) TIPs are tension-responsive proteins involved in myogenic versus adipogenic differentiation. Dev Cell 9: 39–49. doi: 10.1016/j.devcel.2005.04.015

Kingston RE, Chen CA, Rose JK (2003) Calcium phosphate transfection. Curr Protoc Mol Biol Chapter 9: Unit 9 1. doi: 10.1002/0471142727.mb0901s63

Kirino Y, Suzuki T (2005) Human mitochondrial diseases associated with tRNA wobble modification deficiency. RNA Biol 2: 41–4. doi: 10.4161/rna.2.2.1610

Koenig T, Menze BH, Kirchner M, Monigatti F, Parker KC, Patterson T, Steen JJ, Hamprecht FA, Steen H (2008) Robust prediction of the MASCOT score for an improved quality assessment in mass spectrometric proteomics. J Proteome Res 7: 3708–17. doi: 10.1021/pr700859x

Lamichhane TN, Arimbasseri AG, Rijal K, Iben JR, Wei FY, Tomizawa K, Maraia RJ (2016) Lack of tRNA-i6A modification causes mitochondrial-like metabolic deficiency in S. pombe by limiting activity of cytosolic tRNATyr, not mito-tRNA. RNA 22: 583–96. doi: 10.1261/rna.054064.115

Lamichhane TN, Blewett NH, Maraia RJ (2011) Plasticity and diversity of tRNA anticodon determinants of substrate recognition by eukaryotic A37 isopentenyltransferases. RNA 17: 1846–57. doi: 10.1261/rna.2628611

Lentini JM, Alsaif HS, Faqeih E, Alkuraya FS, Fu D (2020) DALRD3 encodes a protein mutated in epileptic encephalopathy that targets arginine tRNAs for 3-methylcytosine modification. Nat Commun 11: 2510. doi: 10.1038/s41467-020-16321-6

Lin H, Miyauchi K, Harada T, Okita R, Takeshita E, Komaki H, Fujioka K, Yagasaki H, Goto YI, Yanaka K, Nakagawa S, Sakaguchi Y, Suzuki T (2018) CO2-sensitive tRNA modification associated with human mitochondrial disease. Nat Commun 9: 1875. doi: 10.1038/s41467-018-04250-4

Mao S, Haruehanroengra P, Ranganathan SV, Shen F, Begley TJ, Sheng J (2021) Base Pairing and Functional Insights into N(3)-Methylcytidine (m(3)C) in RNA. ACS Chem Biol 16: 76–85. doi: 10.1021/acschembio.0c00735

Noma A, Yi S, Katoh T, Takai Y, Suzuki T, Suzuki T (2011) Actin-binding protein ABP140 is a methyltransferase for 3-methylcytidine at position 32 of tRNAs in Saccharomyces cerevisiae. RNA 17: 1111–9. doi: 10.1261/rna.2653411

Olejniczak M, Uhlenbeck OC (2006) tRNA residues that have coevolved with their anticodon to ensure uniform and accurate codon recognition. Biochimie 88: 943–50. doi: 10.1016/j.biochi.2006.06.005

Pan T (2018) Modifications and functional genomics of human transfer RNA. Cell Res 28: 395–404. doi: 10.1038/s41422-018-0013-y

Pereira M, Francisco S, Varanda AS, Santos M, Santos MAS, Soares AR (2018) Impact of tRNA Modifications and tRNA-Modifying Enzymes on Proteostasis and Human Disease. Int J Mol Sci 19. doi: 10.3390/ijms19123738

Phizicky EM, Hopper AK (2015) tRNA processing, modification, and subcellular dynamics: past, present, and future. RNA 21: 483–5. doi: 10.1261/rna.049932.115

Putz J, Dupuis B, Sissler M, Florentz C (2007) Mamit-tRNA, a database of mammalian mitochondrial tRNA primary and secondary structures. RNA 13: 1184–90. doi: 10.1261/rna.588407

Rath S, Sharma R, Gupta R, Ast T, Chan C, Durham TJ, Goodman RP, Grabarek Z, Haas ME, Hung WHW, Joshi PR, Jourdain AA, Kim SH, Kotrys AV, Lam SS, McCoy JG, Meisel JD, Miranda M, Panda A, Patgiri A, Rogers R, Sadre S, Shah H, Skinner OS, To TL, Walker MA, Wang H, Ward PS, Wengrod J, Yuan CC, Calvo SE, Mootha VK (2021) MitoCarta3.0: an updated mitochondrial proteome now with sub-organelle localization and pathway annotations. Nucleic Acids Res 49: D1541–D1547. doi: 10.1093/nar/gkaa1011

Richter U, Evans ME, Clark WC, Marttinen P, Shoubridge EA, Suomalainen A, Wredenberg A, Wedell A, Pan T, Battersby BJ (2018) RNA modification landscape of the human mitochondrial tRNA(Lys) regulates protein synthesis. Nat Commun 9: 3966. doi: 10.1038/s41467-018-06471-z

Sakurai M, Ohtsuki T, Watanabe K (2005) Modification at position 9 with 1-methyladenosine is crucial for structure and function of nematode mitochondrial tRNAs lacking the entire T-arm. Nucleic Acids Res 33: 1653–61. doi: 10.1093/nar/gki309

Schmidt TG, Batz L, Bonet L, Carl U, Holzapfel G, Kiem K, Matulewicz K, Niermeier D, Schuchardt I, Stanar K (2013) Development of the Twin-Strep-tag(R) and its application for purification of recombinant proteins from cell culture supernatants. Protein Expr Purif 92: 54–61. doi: 10.1016/j.pep.2013.08.021

Suzuki T (2021) The expanding world of tRNA modifications and their disease relevance. Nat Rev Mol Cell Biol. doi: 10.1038/s41580-021-00342-0

Suzuki T, Nagao A, Suzuki T (2011) Human mitochondrial tRNAs: biogenesis, function, structural aspects, and diseases. Annu Rev Genet 45: 299–329. doi: 10.1146/annurev-genet-110410-132531

Suzuki T, Suzuki T (2014) A complete landscape of post-transcriptional modifications in mammalian mitochondrial tRNAs. Nucleic Acids Res 42: 7346–57. doi: 10.1093/nar/gku390

Suzuki T, Yashiro Y, Kikuchi I, Ishigami Y, Saito H, Matsuzawa I, Okada S, Mito M, Iwasaki S, Ma D, Zhao X, Asano K, Lin H, Kirino Y, Sakaguchi Y, Suzuki T (2020) Complete chemical structures of human mitochondrial tRNAs. Nat Commun 11: 4269. doi: 10.1038/s41467-020-18068-6

The M, MacCoss MJ, Noble WS, Kall L (2016) Fast and Accurate Protein False Discovery Rates on Large-Scale Proteomics Data Sets with Percolator 3.0. J Am Soc Mass Spectrom 27: 1719–1727. doi: 10.1007/s13361-016-1460-7

Tiranti V, Savoia A, Forti F, D’Apolito MF, Centra M, Rocchi M, Zeviani M (1997) Identification of the gene encoding the human mitochondrial RNA polymerase (h-mtRPOL) by cyberscreening of the Expressed Sequence Tags database. Hum Mol Genet 6: 615–25. doi: 10.1093/hmg/6.4.615

Vare VY, Eruysal ER, Narendran A, Sarachan KL, Agris PF (2017) Chemical and Conformational Diversity of Modified Nucleosides Affects tRNA Structure and Function. Biomolecules 7. doi: 10.3390/biom7010029

Watanabe Y, Suematsu T, Ohtsuki T (2014) Losing the stem-loop structure from metazoan mitochondrial tRNAs and co-evolution of interacting factors. Front Genet 5: 109. doi: 10.3389/fgene.2014.00109

Woodson SA, Koculi E (2009) Analysis of RNA folding by native polyacrylamide gel electrophoresis. Methods Enzymol 469: 189–208. doi: 10.1016/S0076-6879(09)69009-1

Xu L, Liu X, Sheng N, Oo KS, Liang J, Chionh YH, Xu J, Ye F, Gao YG, Dedon PC, Fu XY (2017) Three distinct 3-methylcytidine (m(3)C) methyltransferases modify tRNA and mRNA in mice and humans. J Biol Chem 292: 14695–14703. doi: 10.1074/jbc.M117.798298

Yokogawa T, Watanabe Y, Kumazawa Y, Ueda T, Hirao I, Miura K, Watanabe K (1991) A novel cloverleaf structure found in mammalian mitochondrial tRNA(Ser) (UCN). Nucleic Acids Res 19: 6101–5. doi: 10.1093/nar/19.22.6101

Zhang K, Lentini JM, Prevost CT, Hashem MO, Alkuraya FS, Fu D (2020a) An intellectual disability-associated missense variant in TRMT1 impairs tRNA modification and reconstitution of enzymatic activity. Hum Mutat 41: 600–607. doi: 10.1002/humu.23976

Zhang LH, Zhang XY, Hu T, Chen XY, Li JJ, Raida M, Sun N, Luo Y, Gao X (2020b) The SUMOylated METTL8 Induces R-loop and Tumorigenesis via m3C. iScience 23: 100968. doi: 10.1016/j.isci.2020.100968

